# Mechanically induced topological transition of spectrin regulates its distribution in the mammalian cortex

**DOI:** 10.1101/2023.01.02.522381

**Authors:** Andrea Ghisleni, Mayte Bonilla-Quintana, Michele Crestani, Atsushi Fukuzawa, Padmini Rangamani, Nils Gauthier

## Abstract

The cell cortex is a dynamic assembly that ensures cell integrity during passive deformation or active response by adapting cytoskeleton topologies with poorly understood mechanisms. The spectrin meshwork ensures such adaptation in erythrocytes and neurons. Erythrocytes rely on triangular-like lattices of spectrin tetramers, which in neurons are organized in periodic arrays. We exploited Expansion Microscopy to discover that these two distinct topologies can co-exist in other mammalian cells such as fibroblasts. We show through biophysical measurements and computational modeling that spectrin provides coverage of the cortex and, with the intervention of actomyosin, erythroid-like lattices can dynamically transition into condensates resembling neuron-like periodic arrays fenced by actin stress fibers. Spectrin condensates experience lower mechanical stress and turnover despite displaying an extension close to the contour length of the tetramer. Our study sheds light on the adaptive properties of spectrin, which ensures protection of the cortex by undergoing mechanically induced topological transitions.

## Introduction

The plasma membrane is a responsive composite material where the lipid bilayer and a plethora of attached cortical proteins create a continuously adapting interface with the extracellular environment. Among these attachments, the actin-spectrin meshwork constitutes a ubiquitous non-polarized and self-assembled scaffold critical in preserving cell shape and integrity, extensively studied in red blood cells (RBCs) and neurons [1, 2]. In erythrocytes, the meshwork imposes the peculiar biconcave shape and acts as the unique dissipator of shear forces that RBCs experience in the bloodstream. Pathogenic spectrin variants affect the meshwork integrity and result in hemolytic diseases that underlie the fundamental role in RBCs integrity and protection against mechanical cues [3, 4]. In neuronal axons, spectrin acts as a ruler that spatially organizes critical membrane components such as ion channels and cortical adapters (i.e., ankyrins), while pathological variants are linked to neurodevelopmental and neurodegenerative conditions [5, 6]. The physiological importance of spectrin is also supported by murine knockout models presenting embryonic lethality due to neurological and cardiovascular defects [7].

At structural level, the building block of this meshwork consists of tetrameric head-to-tail (αβ)_2_-spectrin dimers [8]; these rod-shape tetramers are cross-linked by short actin filaments (∼35 nm in length) via a pair of actin binding domains (ABD) present at each end of the β subunits. Hierarchical meshwork organization is obtained by the binding of multiple spectrin tetramers to a unique short actin filament, which functions as a connecting node. Of the different paralogue genes expressed by the human genome (2 α and 5 β), only the pair αI-βI display the restricted expression pattern to RBCs. Pioneering work by electron microscopy unveiled a triangular-like lattice organization of spectrin in erythrocytes [9], where 5-7 spectrin tetramers are connected via a single actin node. Although discrepancies on lattice dimensions were reported depending on sample preparation method, this conformational arrangement long stood as the unique spectrin organization described, until the advent of optical super-resolution microscopy approaches and the identification of a periodic actin-spectrin array of regular spacing (∼180-190 nm in length) along axons [10]. Alternative ultrastructural spectrin topological organizations in other cellular lineages have not been described. Outside of neurons and RBCs, heterogeneous spectrin distribution in epithelial cells and the accumulation at the intercalated disk in muscle tissue have been observed at optical microscopy resolution [11, 12]. We recently reported similar observations in many other cell types [13]. Latest work in erythrocytes using super resolution approaches also suggests a more heterogeneous distribution with discrete organization of nanoscale clusters [14]. Those observations support the possibility that spectrin might create discrete mesoscale cortical domains of different densities (in the order of 1-10 µm^2^ scale) with specific and specialized topological organizations and functions that remain to be elucidated.

Dynamically, key molecular features have been extrapolated from *in vitro* protein mechanics [15, 16]. The unfolding/refolding capacity of spectrin repeats, triple-helix domains arranged in series that constitute the central portion of the rod-shaped spectrin subunits, have led to the conclusion that spectrin and related proteins (i.e., actinin, utrophin, nesprin, etc.) might possess stretch/relaxation capabilities upon mechanical cues also *in vivo*. However, a direct translation of those mechanical capabilities in a cellular context remains elusive and attempts to elucidate these mechanisms have been limited again to the neuronal and erythroid backgrounds. Indeed, a role for spectrin as a shock-absorber in neurons has been described [17], following evidence in nematodes where spectrin stabilizes sensory neurons against mechanical deformation [18]. In erythrocytes, the actin-spectrin meshwork is the only structural cytoskeleton [4]; therefore, this cell system represents the most simplistic model to study the meshwork mechanical properties and its interplay with the plasma membrane. As for the topological organizations, the mechanical context of the meshwork is unexplored in other cellular systems that largely express spectrin. To measure forces sustained by cytoskeletal components and to map the mechanical stress landscape at subcellular level, different genetically encoded FRET-based sensors have been designed. Krieg *et al.* hypothesized a constitutive mechanical tension in the spectrin meshwork in peripheral axons of nematodes, being able to measure a local decrease in stress upon laser axotomy procedure [19]. Orientation-based FRET sensor inserted in αII-spectrin has been developed as a proxy to estimate cortical forces in eukaryotic cells [20, 21]. Given the hypothesized role as a load-carrying spring or tension absorber, one may wonder whether these heterogeneous distributions, topologies and mechanical states of spectrin may act in concert in the more complex and dynamic organization of the cell cortex.

Here, we use Expansion Microscopy and unveil that mammalian fibroblasts can harbor a combination of erythroid- and neuronal-like spectrin topologies. We demonstrated that spectrin can transition dynamically between these two topologies. This transition is driven by actomyosin contractility and organization. Condensed spectrin adopts parallel arrays of ∼180-190 nm tetramers, forming a pattern fenced by actin stress fibers where spectrin is at low tension and low turnover rate. On the contrary low density spectrin displays higher tension and turnover with molecular organization resembling erythroid-like triangular lattices. Dynamic computational modeling integrating topology, tension and actomyosin allowed us to propose a model of the cell cortex where spectrin undergoes topological and mechanical transitions to regulates its distribution under the plasma membrane.

## Results

### Periodic spectrin organization is found in high density condensates between actin stress fibers

We previously reported that spectrin is dynamically organized in mesoscale cortical domains (1-10 µm^2^ in size) complementary to the actin cytoskeleton during cell spreading [13]. Here, we sought to determine spectrin ultrastructural organization in fully adherent mouse embryonic fibroblasts that exhibit a mature cytoskeleton. Diffraction-limited TIRF microscopy (TIRFM) investigations highlighted discrete zonal compartmentation of endogenous βII-spectrin, which formed high fluorescence intensity condensates without colocalization with F-actin (Figure 1A). In particular, these condensates were preferentially found between ventral stress fibers and never observed in cell protrusions. By analyzing fluorescence intensity distribution at the µm^2^ scale, a long-tail distribution of βII-spectrin emerged and was not observed for F-actin (Figure 1B, Supplementary Figure 1A). Indeed, the distribution of F-actin intensity in the TIRFM plane of illumination (100-200 nm above the surface) better fitted with a two-order decay (Table 1), potentially explained by the existence of two distinct F-actin populations: cortical actin and stress fibers directly linked to adhesion complexes. From the distribution of βII-spectrin intensity, we hypothesized that different and discrete zonal organizations might coexist in the cortex of fibroblasts.

**Figure 1.**
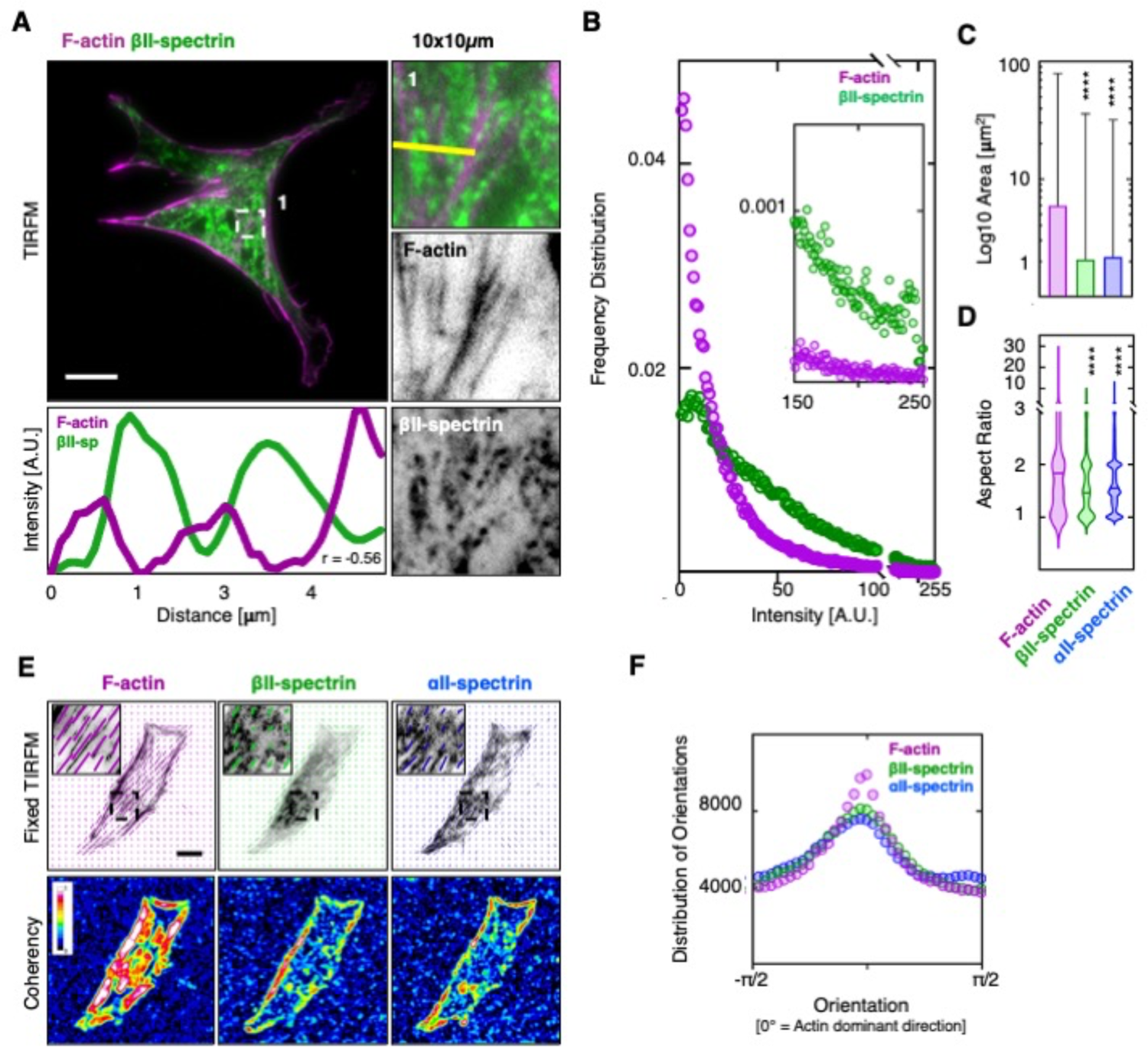
Spectrin condensates align between actin stress fibers. A) MEF immunolabelled for βII-spectrin (green) and F-actin (phalloidin magenta), imaged by total internal reflection microscopy (TIRFM, scale bar = 20 μm). Overlay and single channel images are shown, as well as zooms related to the white dashed box (1). Line scan analysis across the yellow line is reported for both channels to highlight βII-spectrin accumulation between stress fibers (negative Pearson’s correlation coefficient is reported). B) Frequency distribution of F-actin (magenta) and βII-spectrin (green) signal intensities per μm^2^. Inset highlights the different distribution at high intensity ranges (n = 40 cells). Area of high intensity condensates (C) and related aspect ratio (D) highlight smaller and elliptic βII-spectrin and αII-spectrin condensates compared to elongated F-actin (data are presented as mean ± SD, statistical analysis one-way ANOVA with multiple comparisons, **** p< 0.0005). E) Orientation analysis of endogenous βII-spectrin (green), F-actin (phalloidin, magenta) and αII-spectrin (blue). The vectors highlight orientation and coherency of the fluorescent signal in the three independent channels. Coherency heatmaps are shown (LUT 16-colors, scale bar = 20 μm). F) Distribution of orientations of βII-spectrin (green), F-actin (magenta) and αII-spectrin (blue), normalized to F-actin dominant direction (n = 40 cells).

**Table 1:**
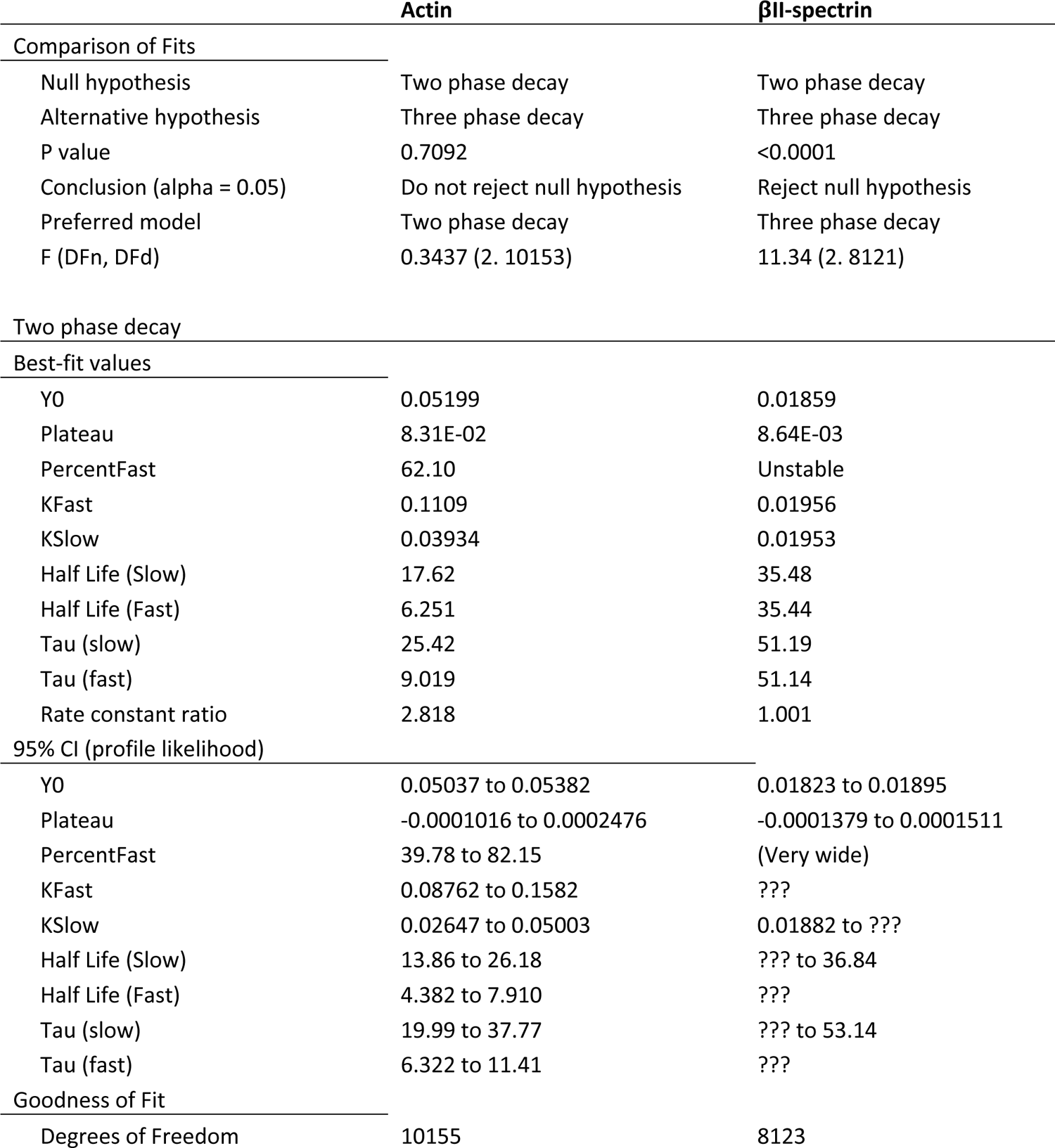
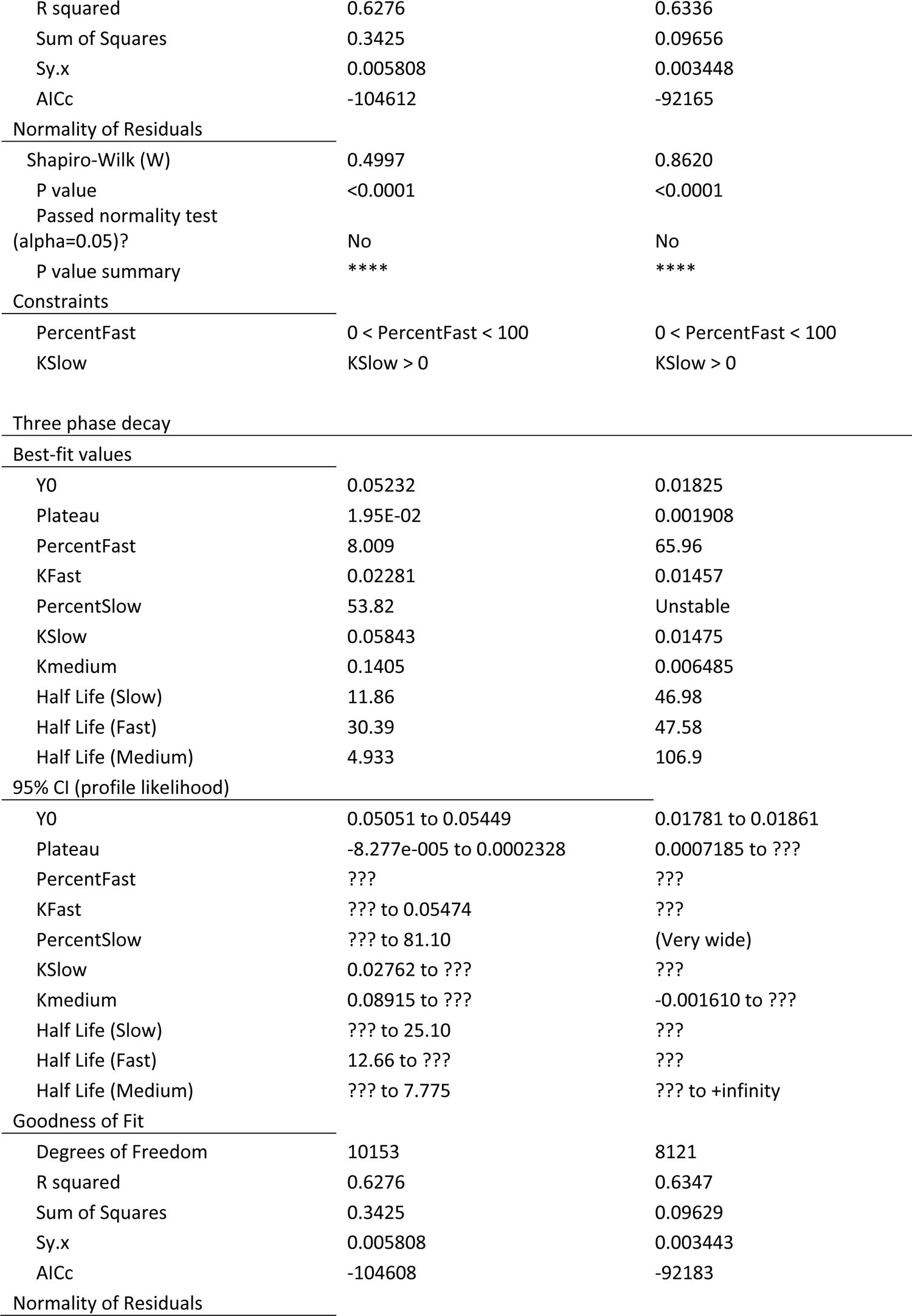
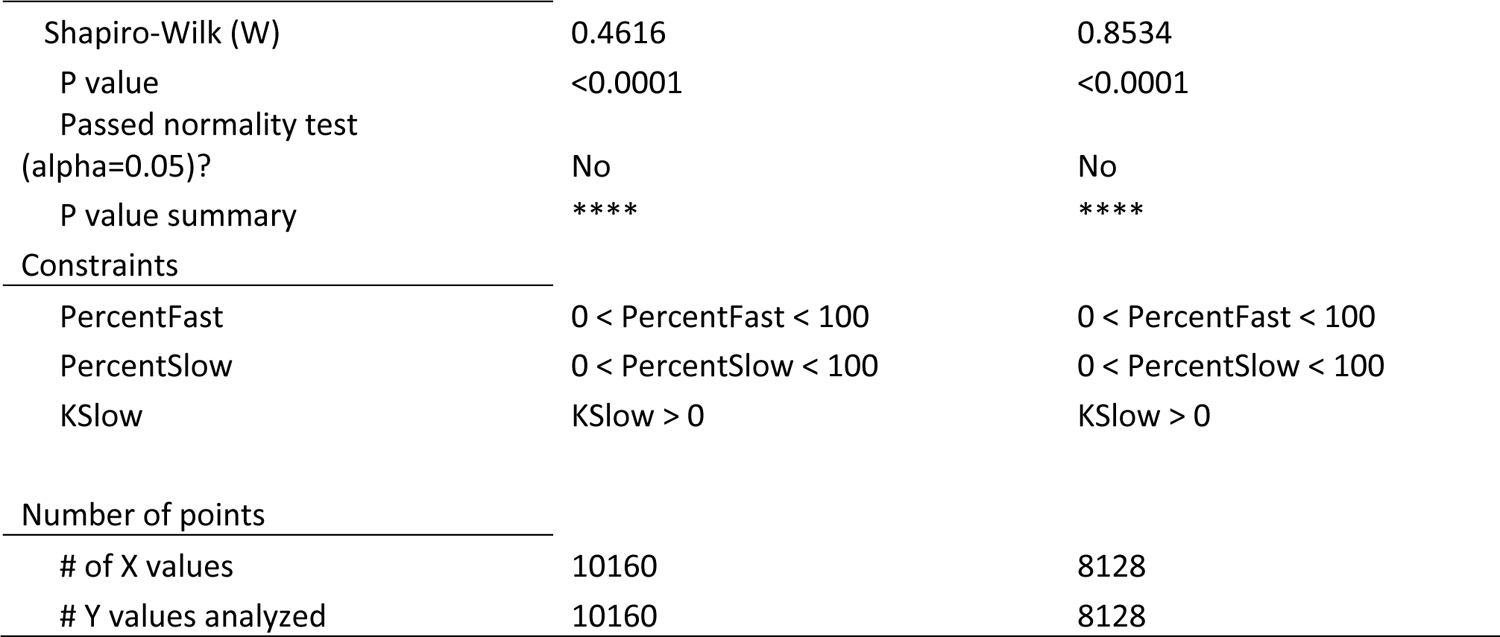
Fluorescence Density analysis

| | Actin | βII-spectrin |
| --- | --- | --- |
| **Comparison of Fits** |  |  |
| Null hypothesis | Two phase decay | Two phase decay |
| Alternative hypothesis | Three phase decay | Three phase decay |
| P value | 0.7092 | <0.0001 |
| Conclusion (alpha = 0.05) | Do not reject null hypothesis | Reject null hypothesis |
| Preferred model | Two phase decay | Three phase decay |
| F (DFn, DFd) | 0.3437 (2. 10153) | 11.34 (2. 8121) |
| **Two phase decay** |  |  |
| **Best-fit values** |  |  |
| Y0 | 0.05199 | 0.01859 |
| Plateau | 8.31E-02 | 8.64E-03 |
| PercentFast | 62.10 | Unstable |
| KFast | 0.1109 | 0.01956 |
| KSlow | 0.03934 | 0.01953 |
| Half Life (Slow) | 17.62 | 35.48 |
| Half Life (Fast) | 6.251 | 35.44 |
| Tau (slow) | 25.42 | 51.19 |
| Tau (fast) | 9.019 | 51.14 |
| Rate constant ratio | 2.818 | 1.001 |
| **95% CI (profile likelihood)** |  |  |
| Y0 | 0.05037 to 0.05382 | 0.01823 to 0.01895 |
| Plateau | -0.0001016 to 0.0002476 | -0.0001379 to 0.0001511 |
| PercentFast | 39.78 to 82.15 | (Very wide) |
| KFast | 0.08762 to 0.1582 | ??? |
| KSlow | 0.02647 to 0.05003 | 0.01882 to ??? |
| Half Life (Slow) | 13.86 to 26.18 | ??? to 36.84 |
| Half Life (Fast) | 4.382 to 7.910 | ??? |
| Tau (slow) | 19.99 to 37.77 | ??? to 53.14 |
| Tau (fast) | 6.322 to 11.41 | ??? |
| **Goodness of Fit** |  |  |
| Degrees of Freedom | 10155 | 8123 |
| R squared | 0.6276 | 0.6336 |
| Sum of Squares | 0.3425 | 0.09656 |
| Sy.x | 0.005808 | 0.003448 |
| AICc | -104612 | -92165 |
| Normality of Residuals |  |  |
| Shapiro-Wilk (W) | 0.4997 | 0.8620 |
| P value | <0.0001 | <0.0001 |
| Passed normality test |  |  |
| (alpha=0.05)? | No | No |
| P value summary | **** | **** |
| Constraints |  |  |
| PercentFast | 0 < PercentFast < 100 | 0 < PercentFast < 100 |
| KSlow | KSlow > 0 | KSlow > 0 |
| Three phase decay |  |  |
| Best-fit values |  |  |
| Y0 | 0.05232 | 0.01825 |
| Plateau | 1.95E-02 | 0.001908 |
| PercentFast | 8.009 | 65.96 |
| KFast | 0.02281 | 0.01457 |
| PercentSlow | 53.82 | Unstable |
| KSlow | 0.05843 | 0.01475 |
| Kmedium | 0.1405 | 0.006485 |
| Half Life (Slow) | 11.86 | 46.98 |
| Half Life (Fast) | 30.39 | 47.58 |
| Half Life (Medium) | 4.933 | 106.9 |
| 95% CI (profile likelihood) |  |  |
| Y0 | 0.05051 to 0.05449 | 0.01781 to 0.01861 |
| Plateau | -8.277e-005 to 0.0002328 | 0.0007185 to ??? |
| PercentFast | ??? | ??? |
| KFast | ??? to 0.05474 | ??? |
| PercentSlow | ??? to 81.10 | (Very wide) |
| KSlow | 0.02762 to ??? | ??? |
| Kmedium | 0.08915 to ??? | -0.001610 to ??? |
| Half Life (Slow) | ??? to 25.10 | ??? |
| Half Life (Fast) | 12.66 to ??? | ??? |
| Half Life (Medium) | ??? to 7.775 | ??? to +infinity |
| Goodness of Fit |  |  |
| Degrees of Freedom | 10153 | 8121 |
| R squared | 0.6276 | 0.6347 |
| Sum of Squares | 0.3425 | 0.09629 |
| Sy.x | 0.005808 | 0.003443 |
| AICc | -104608 | -92183 |
| Normality of Residuals |  |  |
| Shapiro-Wilk (W) | 0.4616 | 0.8534 |
| P value | <0.0001 | <0.0001 |
| Passed normality test<br>(alpha=0.05)? | No | No |
| P value summary | **** | **** |
| Constraints |  |  |
| PercentFast | 0 < PercentFast < 100 | 0 < PercentFast < 100 |
| PercentSlow | 0 < PercentSlow < 100 | 0 < PercentSlow < 100 |
| KSlow | KSlow > 0 | KSlow > 0 |
| Number of points |  |  |
| # of X values | 10160 | 8128 |
| # Y values analyzed | 10160 | 8128 |

Spectrin condensates (defined as P_0.95_ of the intensity distribution, Supplementary Figure 1B) were segmented for both βII-spectrin and αII-spectrin (2.08 ± 34 and 2.18 ± 3 µm^2^ average area respectively), while clusters of bigger size were identified for F-actin (6 ± 73 µm^2^, Figure 1C). Moreover, the aspect ratios (AR) for both spectrins were reminiscent of an elliptic shape (AR: 1.6 ± 0.6 and 1.7 ± 0.7 for βII-spectrin and αII-spectrin), compared to the AR of F-actin which resulted more elongated as expected for long fibers (AR: 2.3 ± 2, Figure 1D). On the same set of images, a matching alignment between F-actin dominant direction with both βII-spectrin and αII-spectrin emerged (Figure 1E and 1F, see Methods for orientation analysis), but with lower level of coherency for spectrins. These results indicated that spectrin condensates have the tendency to align along the direction dictated by F-actin, the cytoskeletal structure with higher coherency.

To gain spatial resolution by immuno-fluorescence, we implemented Expansion Microscopy (ExM) [22]; this approach ensured a consistent ∼4x expansion factor compared to the original size of the biological specimen (Figure 2A-B, Supplementary Figure 2A). Interestingly, βII-spectrin condensates could be observed also by ExM and highlighted a topological organization reminiscent of the periodic pattern identified in axons, but specifically located in the cortical region composed of ventral stress fibers (stained by β-actin antibody, Figure 2C). When epitope inter-distance was calculated (Figure 2D), the values obtained were ∼4x the expected 180-190 nm periodicity originally identified [10] but without the need of additional image deconvolution approaches routinely applied in super-resolution microscopy. As shown in Figure 2 C-D-E, our immuno-fluorescence microscopy investigations support the parallel orientation of the spectrin periodic array to the stress fibers axes. The ∼4x resolution gain obtained by ExM also resulted in a significant reduction in labeling density (Supplementary Figure 2A). As a result, no intermingled actin staining was observed between βII-spectrin puncta, while the conventional phalloidin labeling is known to not resist the gelation and expansion steps of the protocol. We speculated that short actin filaments that connect spectrin tetramers are immunologically secluded in spectrin condensates or do not contain the specific β-actin isoform immunolabelled here. On the other hand, the commercial antibody against αII-spectrin recognizes two specular epitopes on the tetramer that were not resolved by ExM and a clear periodic organization was not obtained for αII-spectrin, neither its localization in respect to the βII-spectrin one (epitopes mapping is shown in Figure 2A). However, dense αII-spectrin regions were found in correspondence of dense βII-spectrin counterparts, and spatial cross-correlation analysis confirmed the *bonafide* of the antibody immunoreactivity (Supplementary Figure 2B). Our results unveiled a non-random organization of spectrin condensates that counteracts the extended one found in the rest of the fibroblast cortex (zooms 1 and 2, Figure 2E). When βII-spectrin Nearest Neighbor Distance (NND) measurements were performed in adjacent regions within the same cell, only a mild decrease was observed between the extended (860 ± 250 nm) and the constrained meshwork configurations (704 ± 192 nm). NND analysis reports the proximity between local maxima. At our resolution nearest epitopes in dense spectrin regions should not be assumed as directly connected as well as the possibility of underestimating the number of maxima for a non-single molecule imaging approach needs to be accounted for. Altogether, these results raise the intriguing possibility that the spectrin meshwork might reversibly transition between different organizations by applying different topological and mechanical constraints (Figure 2G).

**Figure 2.**
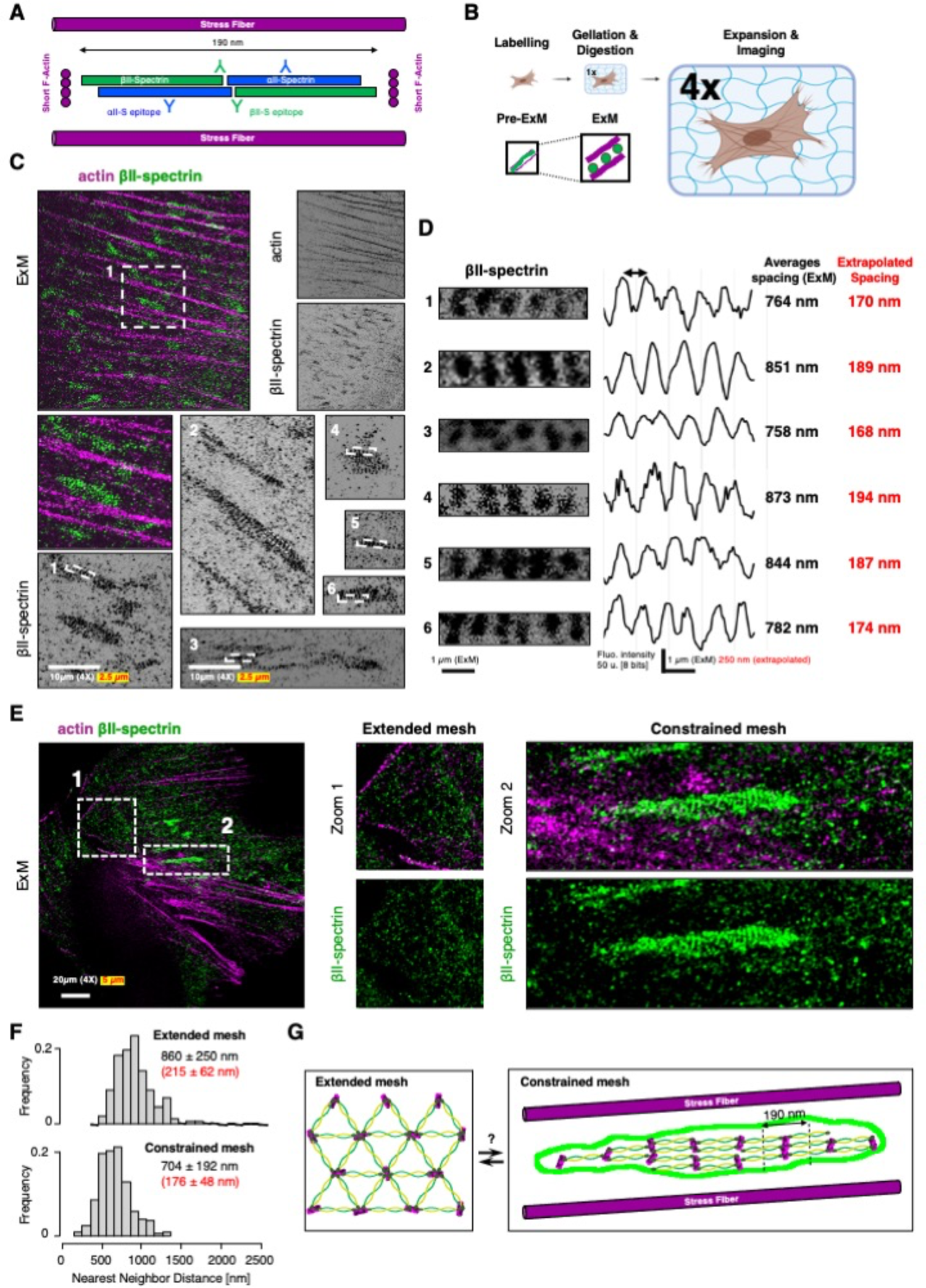
Expansion microscopy unveils the periodic organization of spectrin condensates. A) Cartoon representation of actin-spectrin tetramers (estimated length from the literature ∼180-190 nm) and the relative position of the immuno-reactive epitopes (Y symbols). The experimental pipeline to enhance the spatial resolution by Expansion Microscopy (ExM) is shown in B. C) Representative ExM images of MEF immunolabeled for βII-spectrin (green) and β-actin (magenta) are shown; in white is reported the “real” scale bar of the image (10 μm) and in red the extrapolated scale according to the expansion factor of 4x (2.5 μm). The zooms related to βII-spectrin white box 1 and others (2-6) are shown. Representative images of βII-spectrin periodic condensates are reported in D, and corresponding intensity line scans and peak-to-peak distances (physical in white and extrapolated in red). E) Representative ExM image with two subcellular zones showing βII-spectrin in the extended (zoom 1, white dashed box) and constrained configurations in between stress fibers are shown (zoom 2). F) Nearest Neighbor Distance distributions in the two zooms shown in E are calculated for the βII-spectrin maxima. Adjacent subcellular zones can display dramatic discrepancies in terms of spectrin organizations (drawn in G).

### Actomyosin dynamics controls the formation and disassembly of the spectrin periodic condensates

Next, we asked whether these spectrin periodic condensates were dynamic. Following actin and spectrin by TIRFM, we observed formation and dissipation of GFP-βII-spectrin condensates fenced between stress fibers on the ventral side of the fibroblasts (Figure 3A). As cells change shape and move, condensates follow the framework created by the stress fibers (Figure 3A, zooms 1-6, Movie 1). Those condensates, as well as actin stress fibers, were successfully segmented with the approach previously implemented (Supplementary Figure 1B), and autocorrelation coefficients between consecutive frames were calculated; we observed higher tendency of GFP-βII-spectrin to auto-correlate over time compared to RFP-actin (0.80 ± 0.07 and 0.53 ± 0.12 respectively, Figure 3B-C). While shape descriptors (i.e., size, AR) for those condensates were harder to obtain from live TIRFM datasets, it was possible to analyze the distribution of orientations and relative coherency of GFP-βII-spectrin and RFP-Actin signals. As previously observed for the endogenous staining, GFP-βII-spectrin predominantly aligned with the direction dictated by the more coherent actin cytoskeleton (Figure 3D-E), and sub-cellular regions defined by high βII-spectrin coherency correlated with high actin coherency (Figure 3D, asterisk). Altogether these results confirmed the conclusions previously drawn from fixed TIRFM and ExM investigations corroborating the more passive nature of spectrin condensates compared to the actin cytoskeleton, which is highly dynamic during cell motility. To explore how morphological constraints might regulate spectrin condensation, we engineered the cell culture substrate and seeded fibroblasts on microfabricated fibronectin-coated lines 4 µm in thickness, spaced by 12 µm of non-adhesive surface. On these instructive surfaces, fibroblasts were forced to sprout out thin and elongated processes similar to the proximal portion of axons (Figure 4A-B). ExM on those specimens highlighted the tendency of spectrin to create elongated condensates parallel to the axis of the adhesive lines (Figure 4A-B and Supplementary Figure 3A). However, differently from the axons, βII-spectrin condensates failed to consistently organize into a periodic pattern along the entire length of the cell protrusions, indicating that additional molecular or cell-specific determinants were still required to reach the full barrel and periodic organization described in neurons.

**Figure 3.**
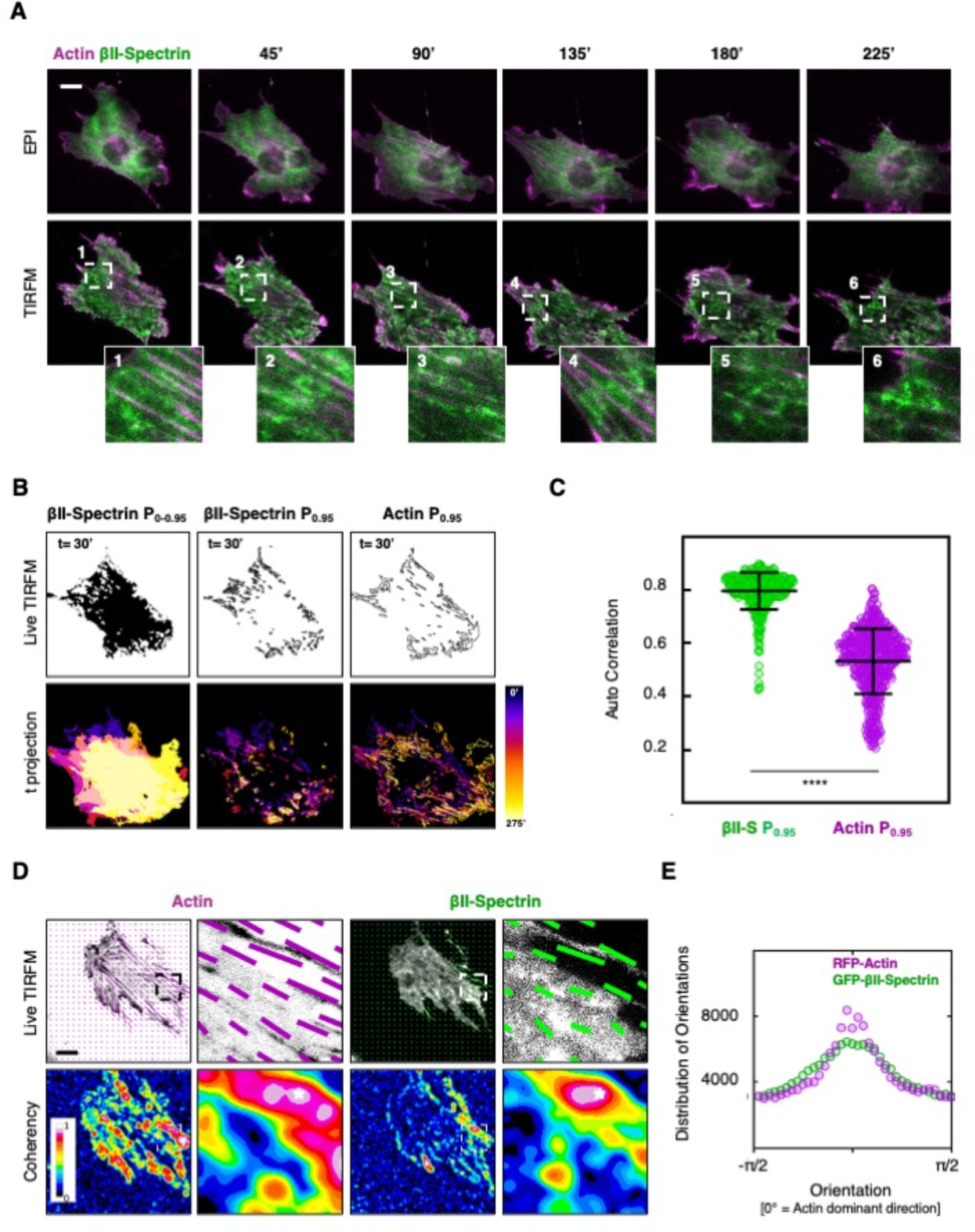
Spatio-temporal evolution of the spectrin condensates. A) Live imaging in dual mode (EPI and TIRF microscopy) of MEF transiently transfected with GFP-βII-spectrin (green) and RFP-actin (magenta). Relevant frames are shown (scale bar = 20 μm), as well as moving zooms of high intensity βII-spectrin condensates only observed by TIRFM (white dashed boxes, 1-6). B) Temporal analysis of GFP-βII-spectrin condensates evolution (P_0.95_ of fluorescent signal intensity distribution is considered). The resulting mask is shown with the condensates outlined at the same time frame (t = 30’), while the color-coded temporal projection is shown for the same cell presented in A, to highlight the dynamic nature of those condensates over time. C) Auto-correlation coefficients between subsequent frames related to GFP-βII-spectrin (green) and RFP-actin (magenta) are shown (n = 8 independent cells and 486 frame pairs analyzed, statistical analysis paired t-test, **** p< 0.0005). D) Representative frames during time lapse analysis of MEF transiently transfected with GFP-βII-spectrin (green) and RFP-actin (magenta). The vectors highlight the orientation and coherency of the signal in the two channels as shown by the heatmaps. E) The graph reports the distribution of orientations of GFP-βII-spectrin (green) and RFP-actin (magenta), normalized to the Actin dominant direction (n = 8 cells in independent time lapses).

**Figure 4.**
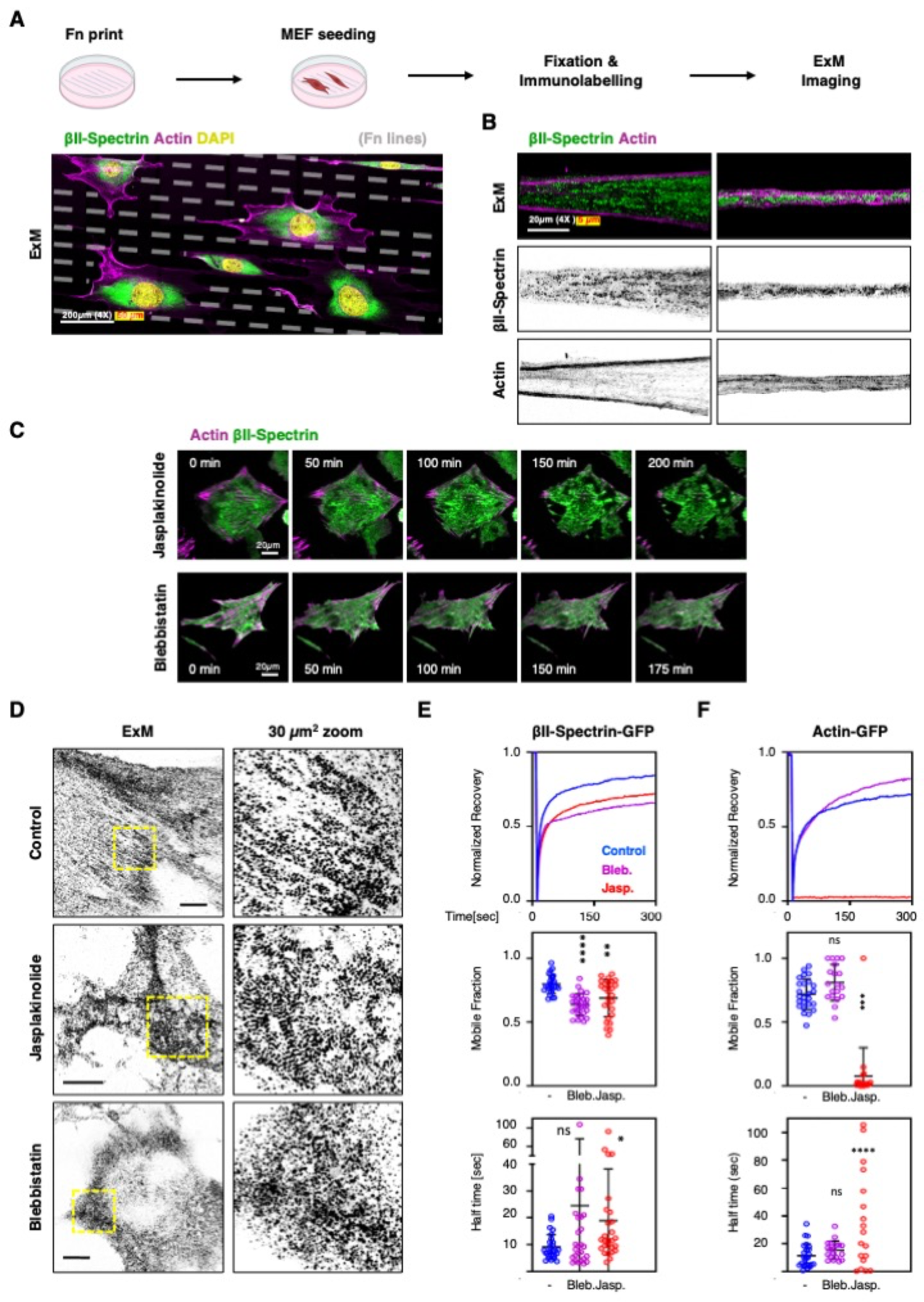
Jasplakinolide treatment drives spectrin condensation. A) Schematic pipeline and representative ExM image of MEFs seeded on microfabricated adhesive lines (4 μm adhesive cross-section, 12 μm non-adhesive surface) and immunolabeled for endogenous βII-spectrin (green) and β-actin (magenta). A large gel volume was imaged at high resolution by tile scan approach, a projection of multiple planes with adjacent cells is shown (scale bar = 200 μm). B) Two different protrusions of independent cells are shown to highlight differential positioning of βII-spectrin and actin (scale bar = 20 μm). C) Live imaging by TIRF microscopy of MEF transiently transfected with GFP-βII-spectrin (green) and RFP-actin (magenta), treated with Jasplakinolide 100 nM and Blebbistatin 10 μM for 4 hours. Relevant frames are shown to highlight the differential effects on cell shape and protein condensation induced by the two drugs (scale bar = 20 μm). D) ExM images of MEFs treated for 3 hours with 10 μM Blebbistatin and 100 nM Jasplakinolide, immunolabeled for βII-spectrin (scale bar = 20 μm). 30 μm^2^ zooms are shown, corresponding to the yellow boxes, to highlight differential effect of the drugs on spectrin organization. E-F) Recovery curves resulting from the FRAP assay are shown for GFP-βII-spectrin and GFP-Actin, transiently transfected in MEFs and treated 3-4 hours with the different cytoskeletal impairing drugs (Jasplakinolide 100 nM, Blebbistatin 10 μM). Mobile fractions are reported in the graphs (statistical analysis one-way ANOVA with multiple comparisons, *** p<0.0005, **** p<0.00005, n = 15-25 independent cells). Half-time recovery rates are reported in the graphs, derived from one-exponential fitting of raw curves (statistical analysis one-way ANOVA with multiple comparisons, * p<0.05, **** p<0.00005, n = 15-25 independent cells).

Pharmaceutical stabilization of F-actin has been previously implemented to obtain enhanced ultrastructural details of the spectrin mesh and actin braids in axons [23]. We treated fibroblasts with 100 nM Jasplakinolide for 3-4 hours in order to stabilize actin cytoskeleton without blocking actomyosin contractility [24]. As a consequence, we observed strong concentration of spectrin signal by formation of super-condensates (Figure 4C, Movie 2). ExM showed that super-condensates were formed by multiple smaller patches of periodic patterns resembling the smaller ones previously observed (Figure 4D and Supplementary Figure 3B). These results suggest that, under Jasplakinolide perturbation, small condensates initially fenced by stress fibers might be dynamically aggregated by myosin contractility. This leads to the formation of spectrin super-condensates made of smaller arrays without dominant orientation, potentially due to the lack of the main directional axis imposed by the stress fibers. When this periodicity followed a clear local directionality, line scan analysis confirmed the expected βII-spectrin periodic spacing of ∼800 nm, the same we previously identified in condensates in untreated fibroblasts (Supplementary Figure 3B). These critical observations support the concept that the spectrin meshwork does not coalesce randomly but instead condense in a controlled manner to form an ultimately packed and ordered lattice of ∼180 nm parallel tetramers.

To gain insights into the molecular mechanisms that can drive the conformational transition of the spectrin meshwork, we measured the effects of Jasplakinolide on protein dynamics by FRAP and observed reduction in mobility and increased half-time recovery for GFP-βII-spectrin in treated fibroblasts compared to untreated cells (Figure 4E, Table 2). The strong negative effect on RFP-actin dynamics was also confirmed with more than 90% reduction in mobility (Figure 4F, Table 2). Interestingly, the effect induced by Jasplakinolide could be reversed and the cells could recover their shape and motility behavior after washout of the compound (Supplementary Figure 4A-B), indicating the reversibility of the process and the retained cell viability despite the strong morphological alterations. On the other hand, myosin-II inhibition by Blebbistatin had opposite effects and promoted homogeneous spectrin distribution without any clear evidence of discrete remaining clusters (Figure 4C-D and Supplementary Figure 3C). As previously reported by us and others [13, 25], Blebbistatin decrease GFP-βII-spectrin dynamics (Figure 4E). Altogether these results support the model that envisions myosin contractility as the main homeostatic driver in spectrin dynamics, required for spectrin condensation into the ultimate periodic organization. This might represent an elegant and non-stochastic mechanoadaptive and mechanoprotective mechanism.

**Table 2:**
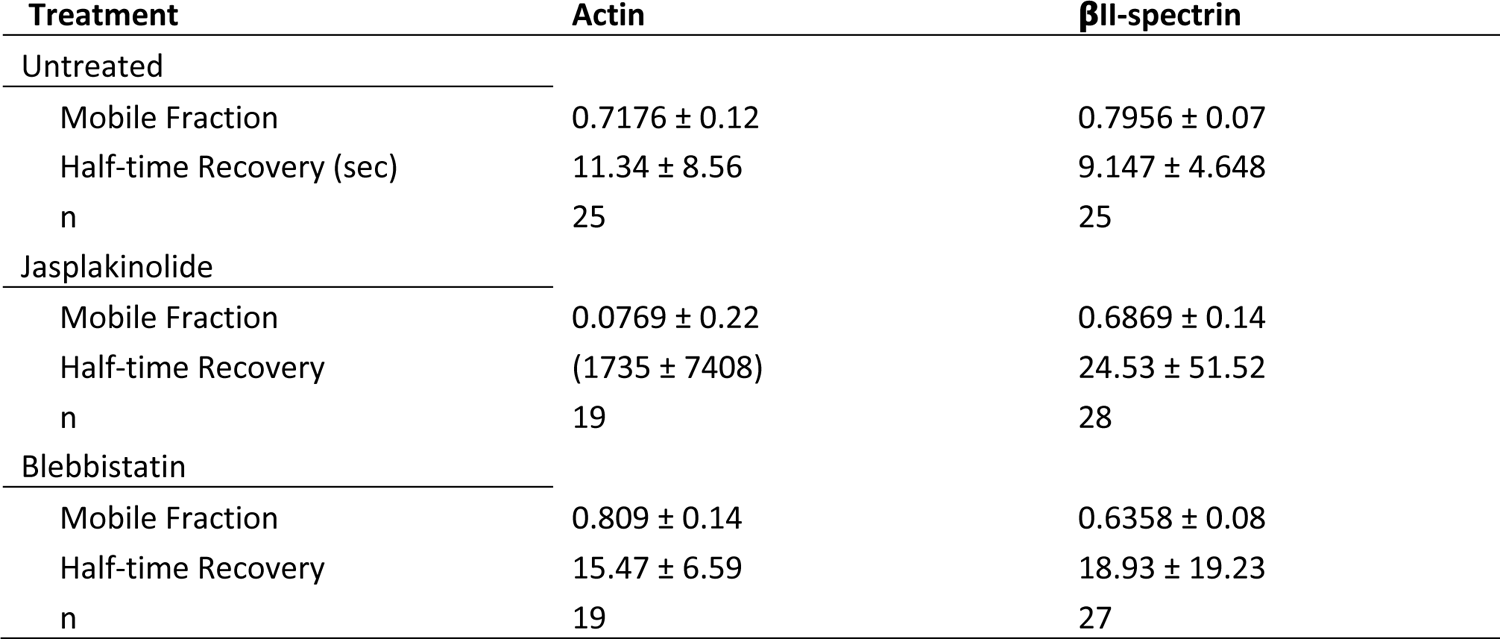
FRAP analysis

| Treatment | Actin | βII-spectrin |
| --- | --- | --- |
| Untreated |  |  |
| Mobile Fraction | 0.7176 ± 0.12 | 0.7956 ± 0.07 |
| Half-time Recovery (sec) | 11.34 ± 8.56 | 9.147 ± 4.648 |
| n | 25 | 25 |
| Jasplakinolide |  |  |
| Mobile Fraction | 0.0769 ± 0.22 | 0.6869 ± 0.14 |
| Half-time Recovery | (1735 ± 7408) | 24.53 ± 51.52 |
| n | 19 | 28 |
| Blebbistatin |  |  |
| Mobile Fraction | 0.809 ± 0.14 | 0.6358 ± 0.08 |
| Half-time Recovery | 15.47 ± 6.59 | 18.93 ± 19.23 |
| n | 19 | 27 |

### Spectrin is in a low tension and low turnover state in the periodic condensates

The spectrin mesh is historically considered as an elastic scaffold that determines the shape of the associated plasma membrane, based on *in vitro* evidences of stretch-and-recoil capabilities upon mechanical stress possessed by the numerous spectrin repeats that form the central rod of the molecule [15, 16]. Considering the results presented here, this molecular feature raises the intriguing possibility that different spectrin organizations might emerge from different tensional landscapes of the cell cortex. To investigate whether this was the case, we envisaged a genetically encoded intramolecular FRET-based βII-spectrin sensor similar to the stress sensor developed for αII-spectrin [21]. Since only the β subunits of the (αβ)_2_ tetramer harbor the actin binding domain (ABD), placing the cpst FRET pair immediately after the “neck” that bridges the ABD with the central rod might provide a more direct strategy to investigate spectrin tensional state compared to the original cpst-αII-spectrin. Moreover, this approach offers the possibility to apply the headless construct lacking the ABD as stress-insensitive reporter (Figure 5A). We measured FRET sensitized emission to derive the inverted FRET ratio as proposed in the original report: lower ratio equals to lower stress and *vice versa* (see Methods). Based on the results presented here, we assumed that spectrin condensates characterized by high fluorescence intensities might present a lower tensional state (low invFRET) compared to regions of the cell characterized by lower fluorescence intensities (measured in the acceptor channel by Venus intensity, Figure 5B). We therefore transiently expressed in fibroblasts cpst-βII-spectrin full length (FL) or the headless ΔABD, and applied the same drug treatments previously described (Figure 4C) to promote different spectrin organizations as means to map the correlation between mechanical stress and local spectrin density. At whole-cell scale, no correlation was identified between the mean invFRET ratio and Acceptor intensity (Supplementary Figure 5A); overall the resulting FRET ratio was not biased by the transfection efficiency of the FRET sensor, which can differ between independent cells and experiments. Moreover, the drug treatments applied did not lead to protein degradation both of the endogenous and the transfected cpst-βII-spectrin FL/ΔABD (Supplementary Figure 5B-C), excluding a contribution of proteolysis in FRET calculations. Semi-automated analysis was performed at μm^2^ resolution for >90 cells per condition (Figure 5C). While all the experimental conditions displayed normal frequency distribution of invFRET values, only in Jasplakinolide treated cells a bimodal distribution emerged, with a specific population of values characterized by low invFRET ratio (low tension). The same population did not appear in the stress-insensitive cpst-βII-spectrin-ΔABD and overall, no significant changes were detected between control and Blebbistatin treatments. We could conclude that when treated by Jasplakinolide, a portion of the spectrin meshwork shifted to a state characterized by lower mechanical stress. To correlate mechanical stress and meshwork density, scatter plots with color-coded Kernel density distributions were generated to visualize the relationship between these two parameters. The population characterized by low invFRET and high fluorescence intensity only appeared when cells expressing cpst-βII-spectrin-FL were treated with Jasplakinolide, and not in all the other experimental conditions tested (Figure 5C, red boxes and bars). Given the observations previously presented by ExM and the FRAP under Jasplakinolide treatment (Figure 4C-D-E, Supplementary Figure 3B and Supplementary Figure 4A-B), we indirectly extrapolated that periodic spectrin condensates corresponded to the regions characterized by lower turnover and lower mechanical stress, and vice versa (see Figure 5D as a cartoon representation of the differential configurations of the spectrin mesh upon drug perturbations).

**Figure 5.**
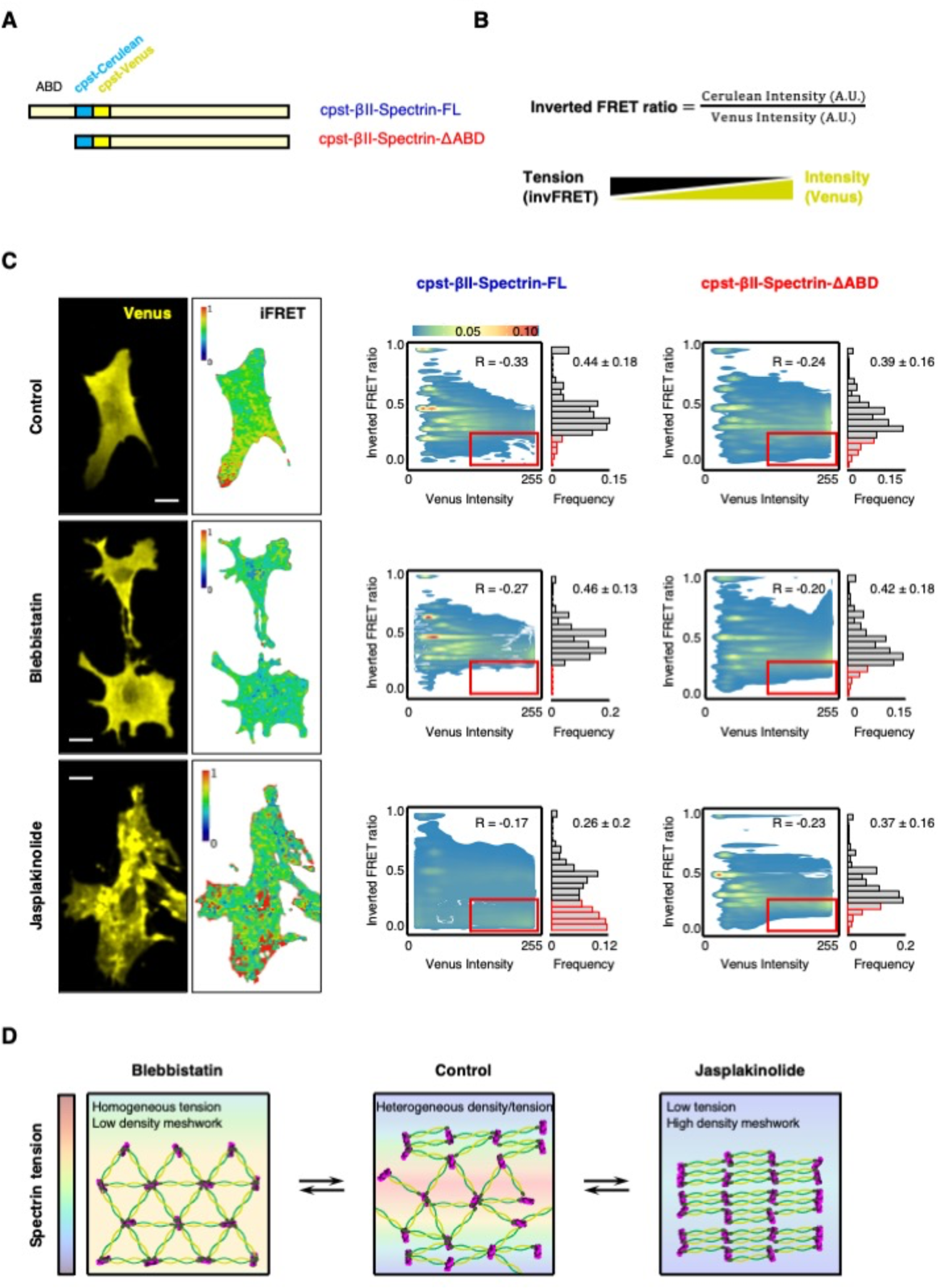
Spectrin presents a low tensional state in condensates. A) Schematic representation of the βII-spectrin FRET-based tension sensor. ΔABD (red) lacks the actin binding domain, while the FRET pair (cpst-Cerulean and cpst-Venus) is inserted between the Actin Binding Domain and the central rod in the full-length counterpart (FL, blue). Inverted FRET ratio is calculated according to Meng and Sachs (2012): higher ratio reflects a higher tensional state (lower FRET) and *vice versa*. The expected relationship between tension and fluorescence intensity of the acceptor (cpst-Venus) is also reported. C) Representative FRET images are shown (scale bar 20 μm). Pixel-by-pixel analysis was performed in at least 90 cells per condition and scatter plots show the local relationship between inverted FRET ratio and Venus intensity. Contours of 2D Kernel densities are reported with the corresponding calibration bar, Pearson’s correlation coefficients are also reported. Frequency distribution of inverted FRET ratio highlight the peculiar behaviour of cpst-βII-spectrin-FL upon Jasplakinolide treatment, which displayed a unique bimodal distribution from all other experimental conditions with the appearance of low inverted FRET ratio values in correspondence of high intensity Venus (highlighted by the red box). Mean ± SD values are reported. D) Cartoon representation of the differential configurations, and relative tensional state (LUT rainbow), of the spectrin meshwork upon drug perturbations.

### Theoretical model indicates that spectrin bundle detachment and active constraint by stress fibers underly conformational changes of spectrin arrays

To investigate how topological constraints can give rise to different spectrin topologies and the related mechanical properties, we developed a computational model. In this mechanical model, the different cytoskeletal elements, namely, filament bundles (tetramers) and crosslinkers are represented by edges and nodes, respectively [26–31]. Spectrin bundles are modeled as edges that behave like springs while short actin filament are the network nodes. By defining the energy generated by the edges and calculating how it affects the position of the nodes (see Methods), we examined the transition between different configurations. First, we considered a network composed of spectrin bundles (edges) arranged in a triangular lattice linked by short actin filaments (nodes). Such a network does mechanical work when the length of its edges is smaller or larger than the resting length of 180 nm derived from experimental observations (Figure 6A, Movie 3). This potential energy exchange produces a restorative force that dictates the movement of the short actin filaments until the edges recover their resting length, and thus, their force is minimized (Figure 6B, Movie 3).

**Figure 6.**
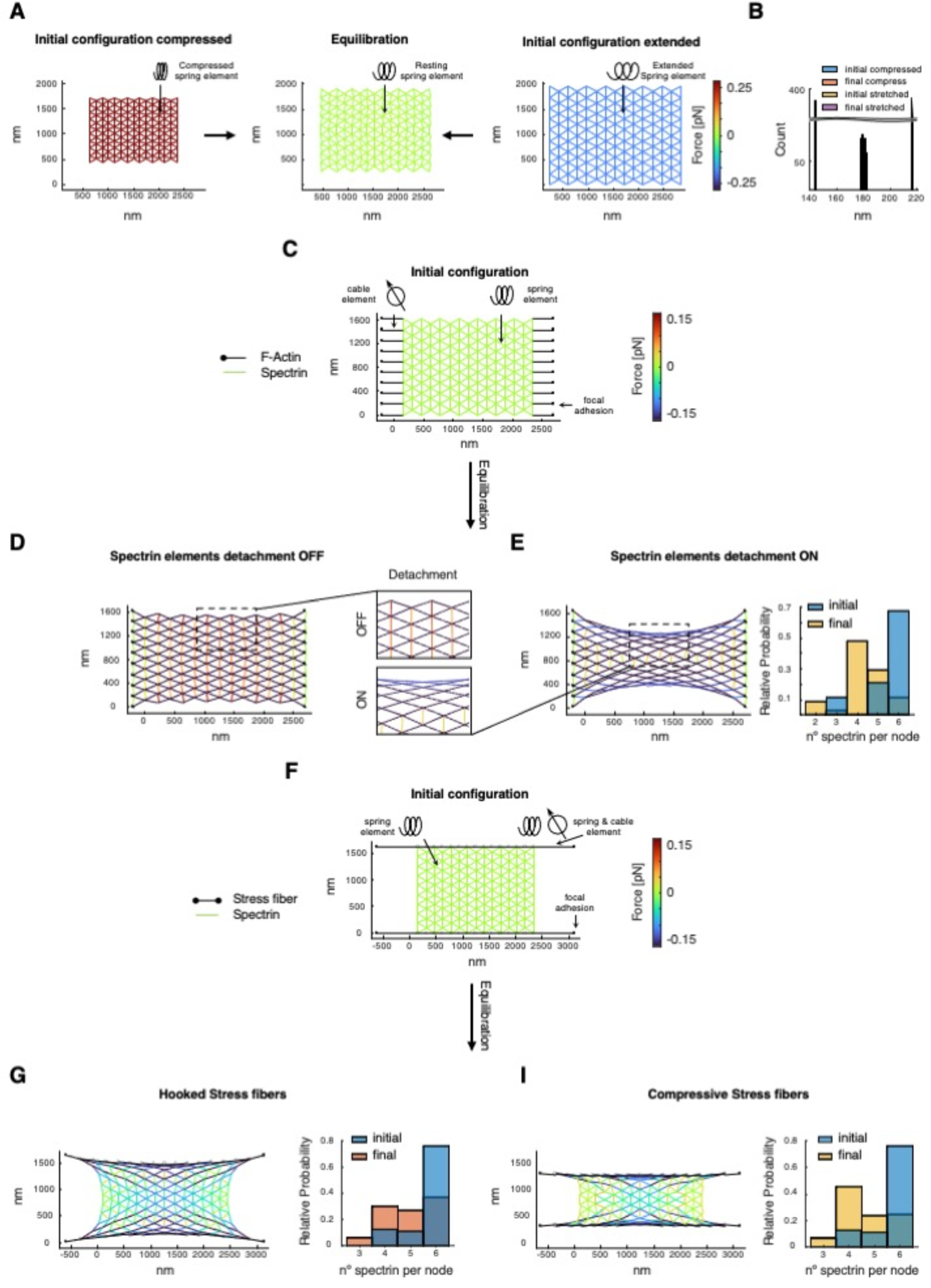
Theoretical model of a spectrin patch. A) Initial configuration of a spectrin mesh (left) where bundles are represented by edges with a spring element, and short-actin filaments are represented by the mesh nodes. The length of each bundle is smaller than the resting length, acting as a compressed spring and exerting a restorative (positive) force on the mesh, depicted by the color of the edges. The configuration of the spectrin mesh in the left evolves to a relaxed state (center), in which the force generated by the spectrin bundles is minimum (equilibration). The time to reach such a state is 60 seconds. Another initial configuration of the spectrin mesh with edges larger than the resting length, acting as stretched springs (right), also relaxes to the configuration shown in the middle. B) Histogram of the initial and final length of the spectrin bundles corresponding to the meshes in (A). C) Spectrin mesh with spring elements attached to fixed focal adhesions (black dots) through cable elements (black edges). D) Final configuration after equilibration (120 seconds) of the network without allowing for any spectrin removal for the mesh shown in (C). E) Configuration of the network in (C) after equilibration allowing spectrin bundles to detach when the force generated by the spring element is above Fth. The histogram shows the number of spectrin bundles per short-actin filament. Zooms of the final configurations are highlighted in squared boxes. F) Spectrin mesh with spring elements attached to stress fibers represented by black edges with spring and cable elements and connected by black empty circles. Solid circles represent fixed focal adhesions. G-I) Final configuration of the network in (F), allowing spectrin bundle removal. In (I), focal adhesions are vertically pulled together during the first half of the simulation. In the second half, focal adhesions are fixed. The total duration is 600 seconds. Histograms represent the number of connected spectrin bundles per short-actin filament.

We hypothesized that when external forces are applied to the network, this triangular erythroid-like configuration changes to a neuronal-like configuration. To test this hypothesis, we considered that the mesh is attached to short cables reminiscent of stress fibers linked to focal adhesion points as seen in cells. Thus, we linked the spectrin mesh to nodes representing focal adhesions through edges with a cable element that exerts a contracting force (Figure 6C). Because we assumed that focal adhesions are anchored to the substrate, the spectrin network stretches (Figure 6D, Movie 4). Note that the resulting network has high mechanical stresses and that the length of spectrin bundles is beyond physiological values. To reduce the mechanical stresses, we introduced spectrin unbinding at a low strain, as suggested in [32], which results in a less stressed configuration (Figure 6E, Movie 4) and the variation of the spectrin bundle length reduces. In such a configuration, the closer the short actin filaments are to the top and bottom of the network, the fewer spectrin bundles they connect, hinting that in the neuronal-like configuration fewer bundles are connected by short actin filaments than the erythroid-like configuration.

Next, we investigated what happens when the spectrin mesh is constrained by stress fibers. To do so, we included stress fibers with a spring and cable element attached to focal adhesions in their ends (Figure 6F). We mimicked the forces generated by actin polymerization in the stress fibers by imposing a greater length in the edge connecting the stress fiber to a focal adhesion [33]. Such a network evolves to a lower-stress configuration (Figure 6G, Movie 5), where the spectrin bundle length and orientation are qualitatively more similar to experimental observations (compare with Figure 1A,E). However, stress fibers are active and are known to move toward each other [34]. We mimicked this motion by moving the focal adhesions toward each other at a constant speed for the first half of the simulation and fixing them thereafter. Figure 6I (and Movie 5) shows that the resulting network relaxes further, and that the movement of stress fibers reduces the number of spectrin bundles connected to a short actin filament.

In conclusion, our model shows that a conformational change in spectrin from erythroid-like to a neuronal-like configuration with lower stress is mediated through the detachment of spectrin bundles with a lower strain, and by active stress fibers. As depicted in Figure 2G, the reduction of the number of spectrin bundles per short actin filament promotes the neuronal-like periodicity in the constrained mesh that otherwise would exhibit a shorter periodicity.

Since our experimental observations pointed to a critical role of myosin contractility in driving spectrin topological transition, we sought to investigate and derive key positional and dynamic parameters from cellular observations to implement myosin workload into this theoretical model.

### Local pulsatile myosin dynamics is predominant in spectrin-rich cortical domains

The contribution of myosin-II-dependent contractility in regulating spectrin dynamics is instrumental for the shape regulation of erythrocytes [25, 35]. We have previously reported reduced spectrin dynamics under Blebbistatin treatment also in fibroblasts [13]. Here we observed that myosin leads to the formation of super-condensates of periodic topology when actin dynamics is impaired (Jasplakinolide treatment). By contrast, blocking myosin contractility while keeping actin dynamics (Blebbistatin treatment) leads to homogenous spectrin distribution without condensation. Therefore, we decided to further investigate the role of myosin-II positioning and contractile outcome in the evolution of cortical spectrin condensates. To visualize myosin structures, we transfected in fibroblasts GFP-tagged myosin-IIA (MIIA) and performed TIRFM investigations. At cortical level, a clear complementarity between spectrin condensates and GFP-MIIA was observed (Supplementary Figure 6A). As expected, MIIA positioning in fixed samples largely matched the F-Actin staining in cells fully adherent to the substrate or confined by the linear fibronectin pattern previously implemented (Figure 4A).

To better understand the spatiotemporal evolution of the actomyosin-spectrin dichotomy, live TIRFM imaging was performed at high temporal resolution (10 seconds frame rate) in fibroblasts transfected for GFP-βII-spectrin and mCherry-myosin light chain (MLC), to unbiasedly label all non-muscle myosin-II isoforms. To emphasize different myosin dynamics depending on different cortical landscapes, fibroblasts were seeded on linear patterns to create controlled discontinuity in cortical territories, either enriched or depleted of stress fibers and adhesions. This strategy allowed us to induce cortical zones of different actin-spectrin densities and myosin-II dynamics (Figure 7A, Movie 6). At single cell level, we observed regions with medium, high and low levels of MLC puncta corresponding to medium, low and high spectrin densities (Figure 7A, zooms 1-2-3, Movie 6). This observation highlighted again the complementarity between actomyosin and the spectrin meshwork, but still lacked dynamic insights. MLC particle tracking was performed and highlighted the appearance of cortical MLC puncta as well as puncta associated with stress fibers (Figure 7B-C, Movie 6). When lifetime of puncta was more carefully analyzed, we observed long-lived MLC tracks to be preferentially associated with the stress fibers (track length > 150 seconds, Figure 7B,E). Instead, short-lived MLC pulses stochastically occurred also in correspondence of denser GFP-βII-spectrin signal (Figure 7A zoom 3, and Figure 7C,E), suggesting a more pulsatile MLC dynamics occurring in cortical regions characterized by prominent spectrin enrichment. These stochastic cortical myosin-II pulses have been previously observed to be dependent on the local recruitment and activation of contractility rather than long-range diffusion of pre-assembled myosin rods [36]. Altogether these results pointed to the possibility that transient local association of myosin-II at regions characterized by the absence of stress fibers is instrumental in driving spectrin condensation, as it is also shown in the kymograph analysis presented (Figure 7D).

**Figure 7.**
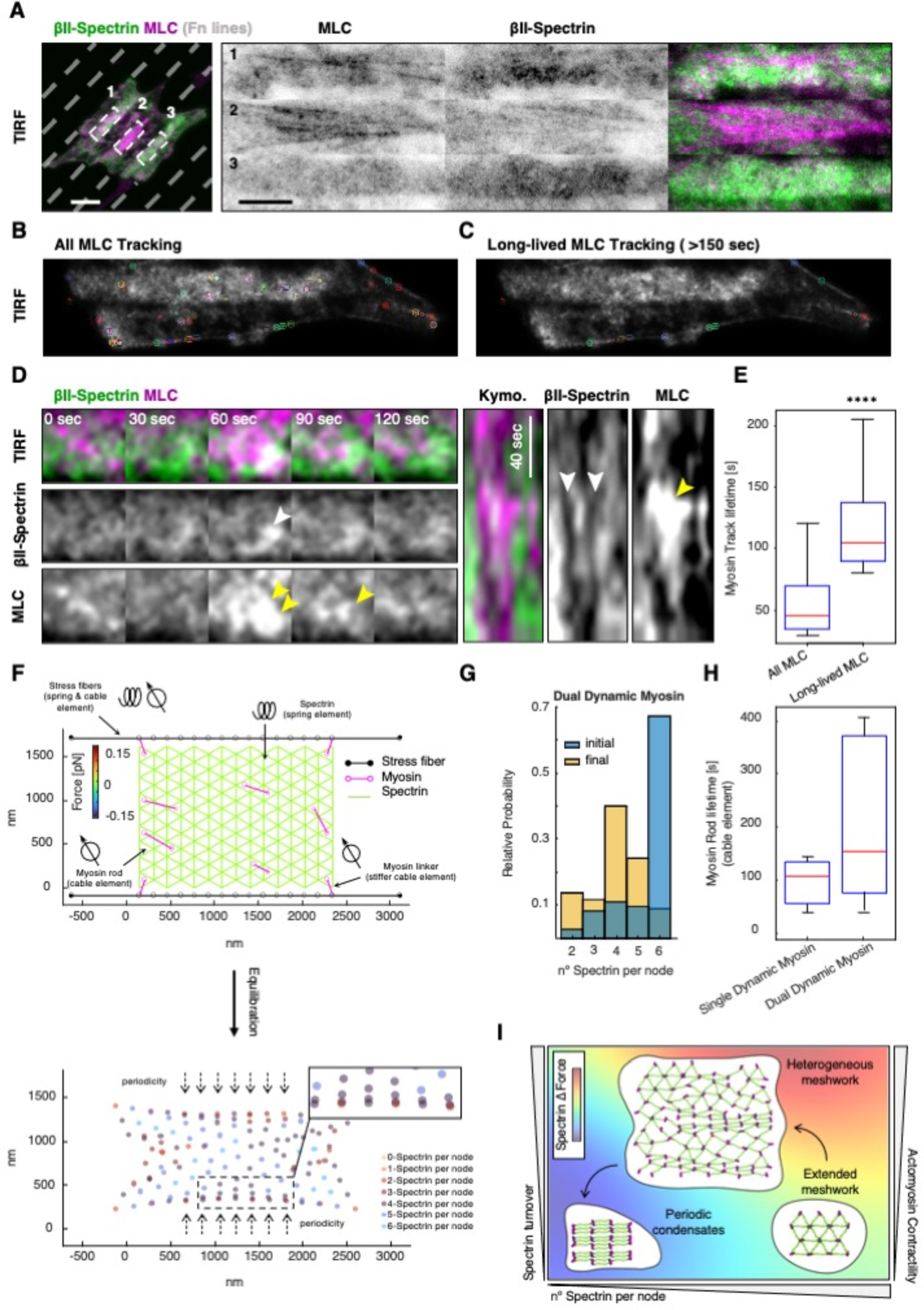
Myosin II dynamics acts as the dominant force for spectrin conformational transitioning. A) Live imaging by TIRF microscopy of MEF transiently transfected with GFP-βII-spectrin (green) and mCherry-MLC (magenta) seeded on microfabricated adhesive lines (not stained, grey) to force discrete cortical organization (scale bar = 20 μm). Three different regions are highlighted and displayed representative cortical dynamics (1-3). Automated tracking of mCherry-MLC puncta with two different filtering parameters: in B, all tracks are shown in a single representative frame, while in C only long-lived tracks (>150 seconds) preferentially localized on stress fibers. D) Pulsative myosin dynamics is shown at spectrin-rich cortical domains (5 x 5 μm zoom). GFP-βII-spectrin and mCherry-MLC condensation is highlighted by white and yellow arrowheads respectively. Kymograph analysis showed fenestration in the GFP-βII-spectrin signal (white arrowheads) in correspondence of mCherry-MLC pulse (yellow arrowhead). E) Differential lifetime of MLC puncta presented in B (All MLC Tracking) and C (Long-lived MLC Tracking). F) Initial configuration of the theoretical model where the spectrin mesh is attached to the stress fibers (black edges) through myosin linkers with cable elements (magenta lines, with empty circles at their ends). Myosin rods with less rigid cable elements are randomly distributed in the mesh. Focal adhesions are pulled together at a constant velocity during the first 300 seconds of the simulation. Afterwards, they are fixed. The total duration of the simulation is 600 seconds. Position of the short-actin filaments at the end of the simulation (equilibration), color-coded to represent the number of attached spectrin bundles per node. Histograms denote the initial and final number of spectrin bundles per short-actin filament (G) and lifetime of the myosin rods in the spectrin network (H) for the two different simulation conditions. I) The phase diagram integrates the different experimental and theoretical parameters identified in this study to explain the contribution of the different players in the topological transition of the spectrin meshwork.

### The inclusion of myosin in the network model results in a more periodic configuration with lower stresses

To examine how myosin affects the spectrin topologies and in agreement with our experimental observations, two different types of myosin dynamics were introduced in the network model: stiff myosin linkers that connect the spectrin mesh to the stress fibers and myosin rods within the spectrin network (Figure 7F, Movie 7), corresponding to the long- and short-lived MLC puncta previously described. Both types of myosin, linkers and rods, are represented by edges with nodes at their ends. Myosins edges are contractile, and we assumed that myosin linkers are stiffer than myosin rods (see Table 3 for modeling parameters). As we never observed clear colocalization between cortical MLC puncta and spectrin, myosin nodes were placed in the center of the triangles formed by spectrin bundles, and when one of these bundles detaches, the corresponding myosin node also detaches and re-attaches to another free spectrin triangle. If there are no free spectrin triangles within a range, then the myosin is removed from the network. Simulations show that the inclusion of myosin alters the final configuration of the network, lowers the stresses, and changes the spatial configuration of the number of spectrin bundles connected by a short actin filament (Figure 7F-G and Supplementary Figure 6B,D,F). Moreover, the final distribution of spectrin length is close to the resting length (Supplementary Figure 6G). We then asked what happens when the myosin rods are added and removed stochastically. In this case, the resulting configuration of the spectrin mesh was improved by reducing the number of spectrin bundles attached to the short actin filaments at the center of the network, thus better resembling the periodic configuration (Figure 7F, Supplementary Figure 6C,E). Furthermore, the model with stochastic myosin rods was able to replicate experimental observations, such as the low number of myosins in the network over time (Figure S6H) and that the dynamic myosin has a shorter lifetime than the myosin attached to the stress fibers, which remains throughout the simulation (Figure 7E and 7H, Movie 7). Note that the contraction generated by myosin allows relaxation of the network in zones that were stressed in the previous model (compare Supplementary Figure 6C,E with Figure 6I). Taken together, our computational model shows that stochastic myosin effectively reduces the stress in the spectrin network and improves its transition to a periodic, neuronal-like configuration.

**Table 3:** Model Parameters

| Parameter | Unit | Value |
| --- | --- | --- |
| $\xi$ : drag coefficient | pN s/nm | 1.25 |
| $\Delta t$ : time step length | s | 0.002 |
| Spectrin |  |  |
| $d_{0,S}$: resting length | nm | 180 |
| $k_{S,S}$: spring constant | pN/nm | 1 |
| $F_{th}$: force threshold for detachment | pN | 0.05 |
| Cable element |  |  |
| $k_c$: cable constant | pN/nm | 150 |
| Stress fibers |  |  |
| $k_{S,F}$: spring constant | pN/nm | 4 |
| $k_{c,F}$: cable constant | pN/nm | 40 |
| $d_{0,F}$: resting length | nm | 165.8846 |
| $t_s$: time for active vertical movement of stress fibers | s | 300 |
| Focal Adhesion |  |  |
| $v_A$: velocity | nm/s | 1.0667 |
| Myosin Linkers |  |  |
| $k_{c,L}$: cable constant | pN/nm | 75 |
| $d_{min}$: minimum length | nm | 135 |
| $d_{max}$: maximum length | nm | 450 |
| Myosin rods |  |  |
| $k_{c,M}$: myosin rod cable constant | nm | 32.1429 |
| $\varphi_a$: myosin rod addition rate | 1/s | 0.01 |
| $\varphi_r$: myosin rod removal rate | 1/s | 0.0063 |

## Discussion

Dynamic changes in cytoskeletal scaffolds are emerging as fundamental mechanisms to convert cell shape adaptation by mechanical cues (mechanoadaptation) into specific activation of transduction pathways (mechanoresponse). Membrane-attached cortical elements represent the primary responsive material that cells can deploy to sense and react to these external perturbations, but are inserted in a crowded milieu notoriously difficult to disentangle by optical and electron microscopy [37, 38]. Here, we identified different topological organizations of the spectrin meshwork and dynamically investigated the contribution of the local mechanical landscape of the cortex in driving this topological transition. To the best of our knowledge, we reported the first observation of the erythroid-like lattice and the neuron-like periodic array of spectrin, coexisting within cells of different lineage specification. Experimental and literature-derived parameters (see Methods) allowed us to build a computational model that integrates actomyosin dynamics, mechanical forces, spectrin turnover, and predicted the molecular events required for the transition between the two ultimate spectrin conformations. In particular, spectrin detachment as a consequence of reduced mechanical stress represents the necessary geometrical requirement to adopt the periodic meshwork organization; the transition is otherwise impossible by applying the sole mechanical stress on the triangular spectrin lattice. If correct, this model describes spectrin capability to respond to local mechanical fluctuations mainly induced by pulsatile actomyosin contractility, and to imprint within the scaffold conformation a cortical mechanical memory [39]. However, this *in silico* observation that represents an intriguing and elegant stress-adaptation mechanism, requires further experimental evidence that is beyond our technical possibilities.

At the whole-cell scale, a functional role for the asymmetric distribution of spectrin has started to emerge during neuronal cell migration and dendrite development in *C. elegans* [40]. The preferential posterior distribution of spectrin, and the predominant Arp2/3 activity at spectrin-depleted leading edges, suggests how spectrin remodeling influences local cortex mechanics by preventing ectopic actin assembly in migrating neural progenitors. We observed a similar exclusion of spectrin from the lamellipodia of spreading fibroblasts, as well as de novo polymerization of branched actin following the displacement of spectrin induced by compressive stimuli [13]. Here, we provide unprecedented details on the different topologies that can be observed in the heterogeneous distribution of the spectrin meshwork within the same cell cortex. During epithelial morphogenesis, asymmetric spectrin distribution is also critical [41]. Indeed, the correct integration of cell shape regulation, cortical mechanics and Hippo signaling effectors is disrupted in the spectrin-deficient Drosophila retina, which lacks the required apical spectrin accumulation to correctly develop the regular array of hexagonal ommatidia [42, 43]. Moreover, dual spectrin/plastin inhibition results in cytokinesis failure due to disorganization and collapse of the equatorial actomyosin network in *C. elegans* [44]. Altogether, these observations point to the capacity of spectrin to create dense buffering reservoirs in cortical regions characterized by low mechanical stress, by adopting the ultimately packed periodic topology. Upon changes in cortical mechanics, these reservoirs can be deployed to secure a continuous but heterogeneous coverage of the plasma membrane.

Historically, the spectrin scaffold is considered as a skeletal integrator [45], capable of organizing the logistic distribution of a plethora of binding partners, cytoskeletal and membrane trafficking events. We previously reported reduced AP2-dependent endocytic events in correspondence of dense spectrin regions induced by osmotic fluctuations [13]; similarly, in neurons clathrin “packets” associate with the spectrin periodic scaffold along the axons, but never fused to the adjacent plasma membrane, suggesting the existence of a membrane fusion barrier in presence of a dense spectrin array [46]. Recent proteomic investigations identified proteins specifically associated with the periodic scaffold and not with the extended meshwork [47]: in particular the structural WD-repeat containing proteins (i.e., coronins), transmembrane channels and receptors (i.e., potassium and sodium ion channels, GPCRs, RTKs), and the signaling molecule CAMK IIβ. Our observations suggest spectrin condensation as a driver for the clustering of transmembrane and membrane-associated proteins also in different cell lineages where spectrin is highly expressed, and potentially explain the integrative role of shear forces and mechanoresponse recently reported in endothelial cells [48–50], with immediate consequences on cell physiology in homeostasis and disease.

The parameters provided here lend themselves to resolving important questions regarding neuronal development. Indeed, the way in which spectrin reaches the stereotypical 3D barrel-like periodic organization in axons represents an informative and specific developmental mechanism. We identified key events required to transition from the extended erythroid-like to a flat periodic spectrin organization. In particular, actomyosin contractility and the constraints imposed by the actin stress fibers are the main drivers of these topological rearrangements. Knowing that membrane mechanics can also drive spectrin reorganization [13], we can reasonably integrate these observations and hypothesize that developing neurons can pack spectrin into the mature barrel with the additional help of constraints imposed by axonal membrane tubulation. This is suggested by our observations in fibroblasts adhered on printed lines, and it would be interesting to study how this topological spectrin transition can be directed in differentiating neurons seeded on these instructive surfaces.

## Acknowledgments

We are grateful to the IFOM imaging facility personnel, in particular D. Parazzoli, Z. Lavagnino, S. Magni and E. Martini for technical support. IFOM cell culture facility personnel. We thank all the members of Gauthier’s, Scita’s, and Maiuri’s groups for helpful discussion. This work was supported by the following funding: an IFOM starting package and an Italian Association for Cancer Research (AIRC) Investigator Grant (IG) 20716 to N.C.G., by H2020-MSCA individual fellowship (796547) and Fondazione Cariplo Young Investigator Grant (2021-1507) to A.G., NIH Grant Number 1RF1DA055668-01 and by an Air Force Office of Scientific Research Grant FA9550-18-1-0051 to P.R.

## Methods

### Cell culture

Immortalized MEFs derived from RPTP α + /+ murine background [51] were grown in complete media composed of DMEM (Lonza) supplemented with 10% Fetal Bovine Serum (FBS South America, Euroclone) and 2mM L-glutamine at 37 °C and 5% CO2. For imaging experiments, MEFs were seeded on borosilicate glass coverslips of 1½ thickness (Corning) or Nunc Glass Base Dishes (Thermo Fischer Scientific) coated with sterile 10 µg ml^−1^ fibronectin (Roche). Supplier and identifier for all reagents are listed in Table 4. Cytoskeletal drug perturbations were performed by supplementing complete media with 10 µM Blebbistatin and 100 nM Jasplakinolide (Merck) for 3-4 hours.

**Table 4:**
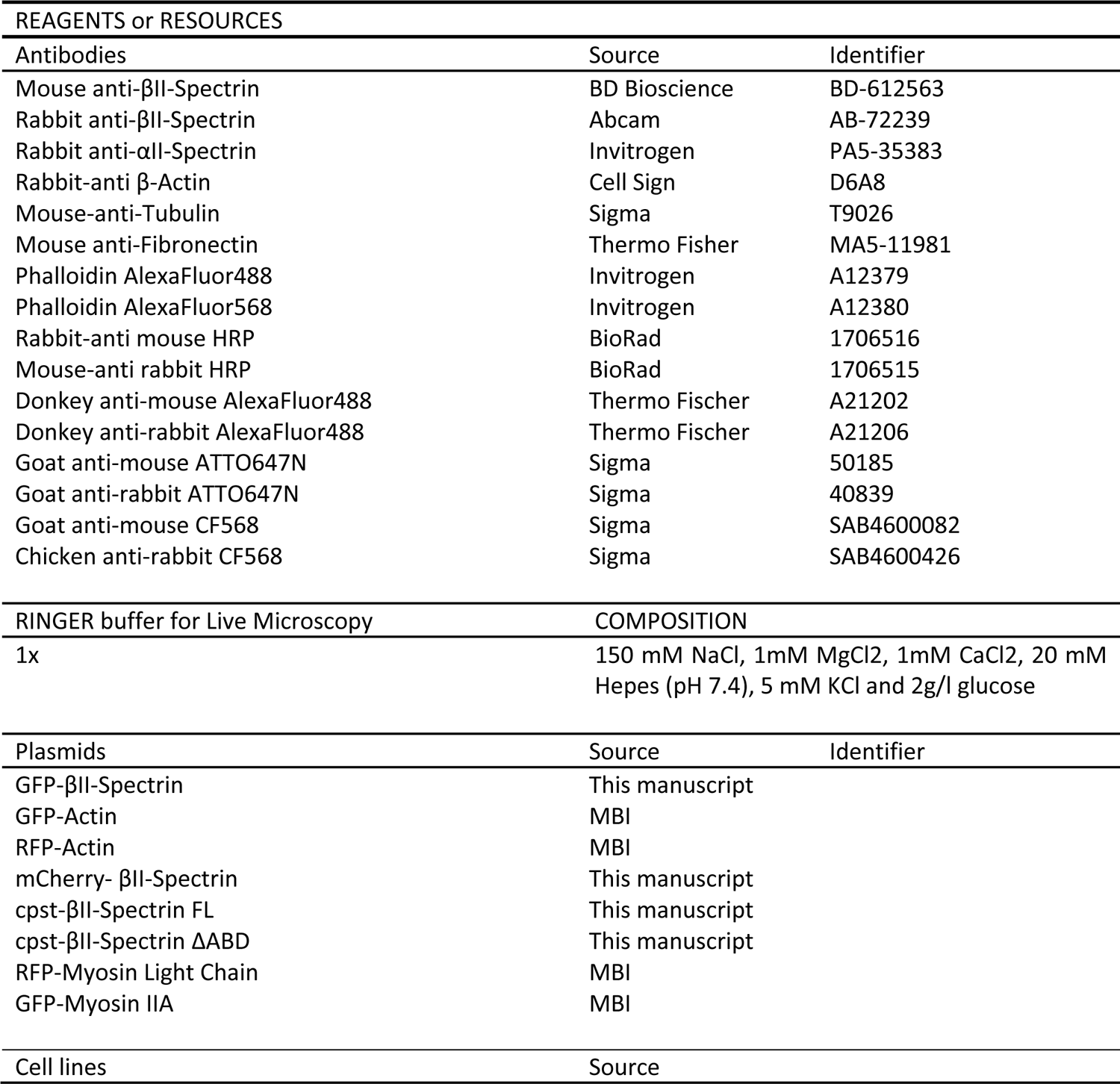
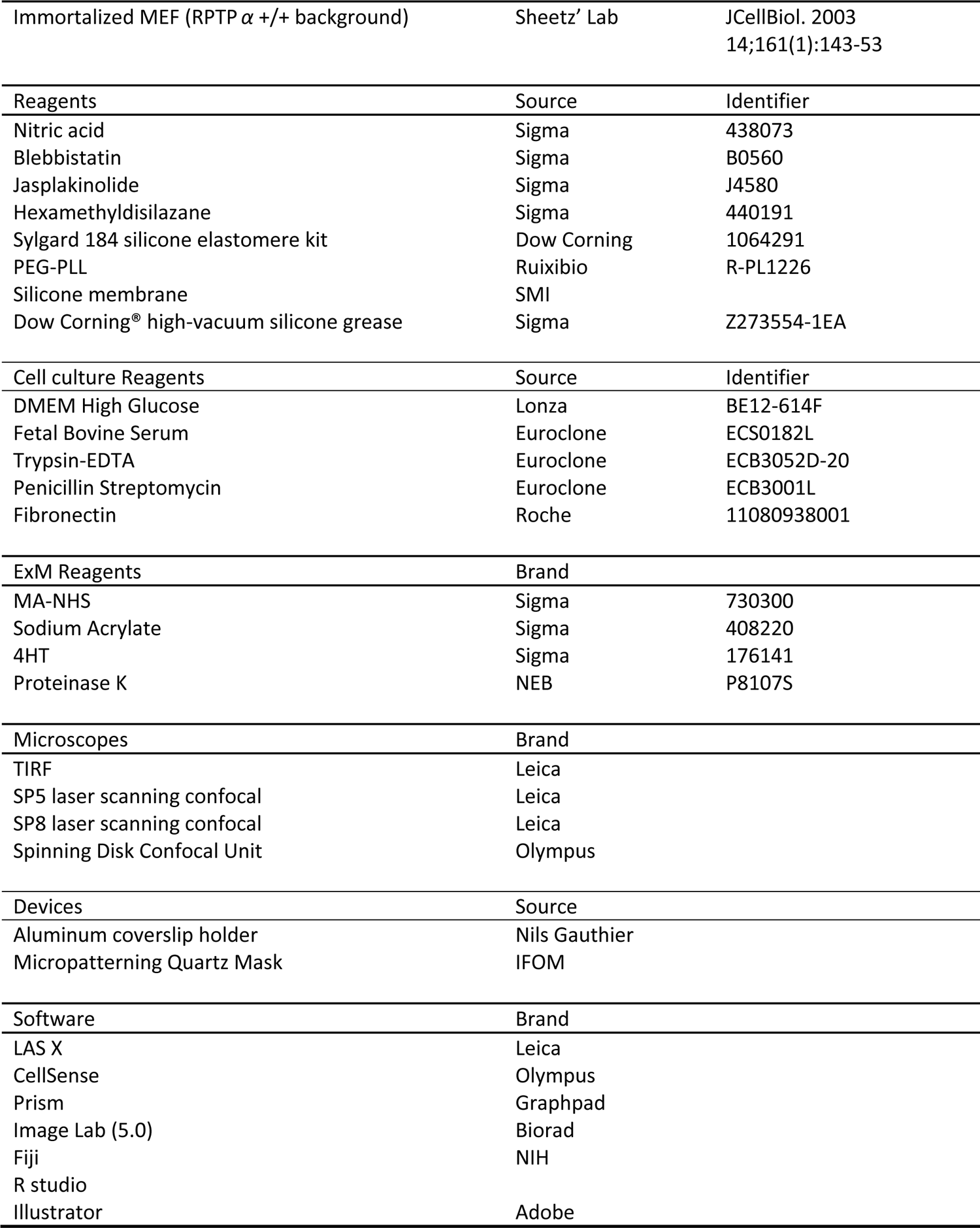
Resources

### Micropatterning

Borosilicate glass coverslips (Corning) were washed for 1 hour with 20% Acetic Acid, rinsed in milliQ water and stored in 90% ethanol. When needed, coverslips were air-dried and activated by a plasma cleaner (Harrick Plasma) for at least 3 minutes. The surface was passivated by incubation with 0.1 mg ml^−1^ poly(ethylene glycol)-b-poly(l-lysine) (PEG–PLL, Ruixibio) for 1 hour at room temperature to prevent fibronectin coating. A quartz mask (Delta mask B.V.) was washed with isopropanol and activated under UV light for 7 minutes (UVO Cleaner, Jelight). PEGylated coverslips were aligned to the desired pattern in the mask, and illuminated under UV light for 7 minutes. The quartz layer prevents UV illumination of the passivated surface while the photolithography-made pattern allows the light to pass, burning the PEG–PLL. Patterned coverslips were then coated with 10 μg ml^-1^ fibronectin for 1 hour at room temperature, while the passivated surface prevented fibronectin adherence. After rinsing the coverslips several times with PBS, MEFs were seeded at desired cell density and cultured at 37°C in the same media described before.

### Immunofluorescence

The antibodies used in this study were the following: mouse anti-βII-spectrin (dilution 1:200, BD-bioscience), rabbit anti-βII-spectrin (1:200, Abcam), mouse anti-αII-spectrin (1:200, Invitrogen), and rabbit anti-β-actin (1:100, Cell Signaling). Before fixation, cells were seeded on 10 μg ml^−1^ fibronectin-coated coverslips/glass base dishes. Cells were fixed in 4% paraformaldehyde for 10 minutes, then neutralized using 10 mM NH_4_Cl in PBS for 10 minutes. Alternatively, fixation was performed in ice-cold pure methanol for 2 minutes at −20 °C. Cells were subsequently washed three times with PBS, permeabilized for 2–5 minutes using PBS containing 0.1% Triton X-100, and blocked with 3–5% BSA for 10 minutes at room temperature. Cells were incubated with primary antibody overnight at 4 °C. After 3 washing steps in PBS, cells were incubated with CF568/AttoN647-conjugated goat anti-mouse or anti-rabbit (1:100–1:400, Thermo Fischer Scientific) and Alexa 488/568-conjugated phalloidin (1:200, Sigma-Aldrich) for 1 hour at room temperature. After three washes in PBS, cells were mounted with anti-fade glycerol-based media (for confocal microscopy) or PBS (for TIRFM and ExM) and stored at 4 °C. All primary antibodies and fluorophore-conjugated secondary antibodies are listed in Table 4.

### Expansion Microscopy (ExM)

The procedure for ExM was adapted from the original report [22]. For completeness of information and reproducibility all sensible steps are described. Immunofluorescence procedure was performed as previously described, except for the fluorophores used: CF568-conjugated goat anti-mouse (1:100 Sigma-Aldrich) and AttoN647-conjugated goat anti-rabbit (1:100, Sigma-Aldrich). When ExM was performed on micropatterned coverslips, before the immunostaining procedure a quenching step of 2 minutes at −20° with ice-cold methanol was required to prevent reactivity of the PEG-PLL surface coating with the amino-reactive Anchoring buffer.

### Anchoring

Upon completion of immunostaining procedure, the specimen on coverslips were incubated with the Anchoring buffer, consisting of PBS supplemented with 1mM of the amino-reactive MA-NHS (Methacrylic Acid N-Hydroxysuccinimide Ester, Sigma-Aldrich). Specimens were incubated for 1-1.5 hours on gentle rocking at room temperature.

### Gelation

Gelation solution was composed of the Monomer stock (sodium acrylate 33 wt%, acrylamide 50 wt%, bis-acrylamide 1 wt%, 1.8M NaCl, 1x PBS, Sigma-Aldrich), supplemented before gelation with 0.2 wt% of TEMED and Ammonium Persulfate (APS, Sigma-Aldrich). Gelation solution was supplemented with 0.01 wt% of 4-Hydroxy-TEMPO (Sigma-Aldrich) to slow down the reaction and allow complete diffusion inside the cells. Anchoring buffer and excess of MA-NHS was removed by 2x brief washes with PBS. Gelation was performed in custom-made gelation chamber: 2 microscopy slides were coated by parafilm and spaced by silicone isolator gaskets of 0.5 mm thickness. Coverslips were placed on the bottom side of the chambers, biological specimens facing the inner side of the chamber, and excess of PBS buffer was removed. Complete Gelation Solution was added and chambers were placed flat on ice for 30 minutes to favor homogeneous diffusion of acrylamide and avoid gelation artefacts. After this step, chambers were placed for 1 hour at 37°.

### Digestion

Gelation chambers were carefully dismounted by removing the top slide and the gel trimmed with a razorblade. Gel shapes with no-mirror symmetry are useful to recognize the surface of the gel with the biological specimens located at the interface. Coverslips with trimmed gels still attached were placed in opportune plastic dish and incubated with Digestion Buffer (50mM Tris-HCl, 125 mM NaCl, 2.5 mM EDTA, 0.5% Triton X-100) supplemented with Proteinase K (1:100, New England Biolabs). Specimens were incubated on an orbital shaker at 37°C per 2 hours and 60 rpm.

### Expansion

For the physical expansion step, gels were transferred into bigger plastic dishes to ensure undisturbed planar expansion. Digestion buffer was replaced by milliQ water and incubated with gentle shacking. At least 4 washing steps of 30 minutes were performed. Expanded gels were kept at 4°C in the dark.

### Mounting

Acrylamide hydrogels required immobilization on glass coverslips to avoid drift during volumetric imaging. Coverslips were plasma activated (Harrick Plasma) for 3 minutes and then coated with poly-L-Lysine for 1 hour at room temperature, washed in milliQ water and dried before letting the gel settle.

### TIRFM and Confocal Microscopy

Confocal microscopy was performed on a Leica TCS SP8 laser-scanning confocal module mounted on a Leica DMi 8 inverted microscope, equipped with a motorized stage, and controlled by the software Leica Application Suite X (ver. 3.5.2.18963). For image acquisition, a HC PL APO CS2 63 × /1.40 oil immersion objective was used. DIC, epifluorescence (EPI), and total internal reflection fluorescence microscopy (TIRFM) of fixed specimens, live time lapses and drug treatments were performed on a Leica AM TIRF MC system. Two different TIRFM-grade objectives were used: HCX PL APO 63 × /1.47NA oil immersion and HCX PL APO 100 × /1.47NA oil immersion. Three different laser lines were used for fluorochrome excitation: 488 nm, 561 nm, and 635 nm. A specific dichroic and emission filter set for each wavelength have been used. The microscope was controlled by Leica Application Suite AF software (ver. 2.6.1.7314), and images were acquired with an Andor iXon DU-8285_VP camera. For live imaging experiments, environmental conditions were maintained by an Okolab temperature and CO_2_ control system.

### Fluorescence Intensity Distribution

TIRFM dataset were analyzed to extract Intensity distribution in Fiji. Briefly, images were acquired at constant laser intensity, exposure time and no electronic gain between independent channels. Images were converted to 8-bit format, and were down-sampled to the pixel size of 1 x 1-µm by interpolating the bilinear average. Binary cell mask was generated by the “Analyze Particle” tool, cell outline was added to the ROI manager. For each channel the Histogram of intensity distribution was generated, the count value matched the projected cell area in µm^2^. Extracted values were normalized by the total count to obtain a relative frequency and allow averaging between multiple cells.

### Condensates segmentation

TIRFM images were processed by the tool “Subtract Background” in Fiji, the method Sliding paraboloid with a rolling ball radius of 50.0 pixels was applied. Single channel images of 16-bit format were independently thresholded with default method, by imposing a cutoff value of 5%. This approach set the 95% of the intensity distribution curve to 0, while the remaining 5% (P_0.95_) is set to max. The Despeckle and Smooth tools were run to homogenize the signal, and the Analyze Particle was used to obtain Area and Shape descriptors for each individual cluster.

### Orientation analysis

To analyze signal orientation on TIRFM dataset the Fiji plugin OrientationJ was used to quantify the local orientation properties of an image, based on the structure tensor of a defined local neighborhood [52, 53]. Briefly, images were background subtracted by the Sliding paraboloid method with rolling ball radius of 50.0 pixels. Vector field was independently calculated for each channel by applying a Gaussian gradient of 5 pixels. Coherency maps as well as overlaid vector images were generated. Distribution of orientations were independently calculated for each channel by the Distribution tool of the plugin. The same Gaussian gradient of 5 pixels was applied, with no Min coherency (0%) set as default. Dominant Direction was instead calculated only for the actin channel, the resulting value for the single cell under investigation was set as 0°. All the other channels were aligned according to actin dominant direction.

### Colocalization analysis

The Fiji plugin JACoP (Just Another Co-localization Plugin) was used to determine co-localization parameters between channels, those includes Pearson’s correlation coefficient and Cross Correlation coefficient with 50-pixel shift. In particular this function returns a series of Correlation coefficients when the two channels are shifted by ± 50 pixels between each other. Raw data were then plotted by using the software Graphpad Prism.

### FRAP experiments

MEFs expressing GFP-βII-spectrin constructs were imaged 24 hours after transfection with a confocal spinning disk microscope (Olympus) equipped with iXon 897 Ultra camera (ANDOR) and a FRAP module furnished with a 405-nm laser. The environmental control was maintained with an OKOlab incubator. Images were acquired using a 100 × /1.35Sil silicone oil immersion objective. MEFs were trypsinized and seeded on glass base dishes (Matek, Sigma-Aldrich) coated with 10 µg ml^−1^ fibronectin. Before imaging, CO_2_-independent 1× Ringer’s solution was exchanged. Circular regions of interest of 3-5 µm diameter were photobleached with the 405-nm laser at 100% intensity, and post-bleach images were acquired with 15–20% laser intensity for 100 frames (1 frame every 3 seconds for full-length and truncated GFP-βII-spectrin constructs and every 0.5 seconds for PE/ANKbs only). FRAP data were analyzed, and curves fitted to the one-exponential recovery equations (one-phase association) by the software Graphpad Prism:

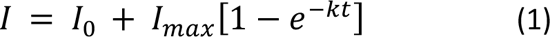

where I represents the relative intensity compared to the prebleach value, k the association rate, and the half-time recovery expressed in seconds.

### Sensitized FRET emission

#### Imaging settings

FRET recordings were performed on cells transfected with cpst-βII-spectrin-FL or cpst-βII-spectrin-ΔABD imaged through a Leica SP8 Confocal microscope. Briefly, three channels were sequentially recorded by an HyD detector with no electronic gain: donor, FRET and acceptor channels. Donor excitation was achieved with an Argon laser at 458 wavelength, and emission filters were set at 475-485nm; FRET excitation with the same 458nm laser and emission filters 515-525nm; acceptor excitation with 514nm laser and emission filters 520-600nm. Images were acquired with pinhole set to 1 AU in 8-bit and 512 x 512-pixel format.

#### Inverted FRET index calculation

Raw images were processed by applying a Gaussian blur of sigma radius of 1 µm, ensuring this spatial resolution in the resulting FRET values. A binary mask of the cell is created and used to set all the values outside the cell to NaN. According to the original report [21], inverted FRET ratio was calculated by dividing the Donor image by the FRET image. Floating values were normalized between 0 and 1. For whole-cell inverted FRET calculations the Analyze particle tool was used to get the Mean grey values. Since outside the cell mask pixels were set to NaN, mean grey values only resulted from the cell projected area. For pixel-by-pixel analysis a stack of 2 channel was generated: the first channel represented the inverted FRET ratio image (mean grey values between 0 and 1) and the second channel the Acceptor image (mean grey value between 0 and 255, 8-bit format). A custom-written plugin allowed the retrieval of values in both channels for corresponding pixels, creating a .csv file to be analyzed in R. Scatter plots at each experimental condition were generated by excluding saturated pixels from the analysis.

#### Western blotting

For western blot analysis, cells were lysed directly on the plate by adding opportune amount of modified Laemmli sample buffer composed of Tris-HCl 135 mM (pH 6.8), sodium dodecyl sulfate (SDS) 5%, urea 3.5 M, NP-40 2.3%, β-mercaptoethanol 4.5%, glycerol 4%, and traces of bromophenol blue. Total protein content was normalized by seeding cells at equal densities; this lysis buffer does not allow total protein quantification, but prevents membrane-bound proteins from being degraded during trypsinization. Equal volume between samples was then loaded onto 12–8% SDS polyacrylamide gels and transferred after electrophoretic separation onto nitrocellulose membrane (Amersham GE-Healthcare). After the transfer, membranes were blocked in PBS supplemented with 0.1–0.3% Tween20 and 5% milk for 1 hour at room temperature, then incubated overnight at 4° with primary antibodies at the following dilutions: mouse anti-βII-spectrin 1:2000 (BD-bioscience), rabbit anti-βII-spectrin 1:2000 (Abcam), and mouse anti-β-tubulin 1:5000 (Sigma-Aldrich). After three washing steps in PBS–Tween20 (0.1–0.3%), membranes were incubated for 1 hour at room temperature with HRP-conjugated secondary antibodies (BioRad). Three washing steps of 5 minutes in PBS–Tween20 (0.1–0.3%) were performed between primary and secondary antibody incubation. Proteins were detected by ECL Western blotting reagents (Amersham GE-Healthcare), using the digital Chemidoc XRS + system run by the software Image Lab (Biorad).

#### Model

We proposed a model to investigate whether the spectrin periodic pattern seen between stress fibers in the cell emerges from a periodic hexagonal pattern, like that of the red blood cell. We hypothesized that this change in the cytoskeleton configuration is possible due to the interaction of the forces generated by components. Therefore, to investigate the mechanical forces of the cell cytoskeleton, we modeled it as a network of springs and cables, as in[26–31], where the edges and nodes correspond to filament bundles and cross-linkers, respectively. This mesoscopic description gives a good approximation for the spatio-temporal evolution of the cytoskeletal components under investigation. Unlike molecular dynamics and dissipative particle dynamics models [25, 32–34], this approach allows the exploration of a larger cytoskeletal meshwork without computationally expensive simulations. Furthermore, we used a 2D network assuming that the out-of-plane deformations are negligible in the cells under consideration.

In the initial model, spectrin bundles were represented by the edges of a triangular mesh connected by short actin filaments (nodes). Such a configuration resembles the cytoskeleton of a red blood cell. Spectrin bundles behave like springs, in the sense that they return to a resting length after stretching or shrinking. Hence, the N*_e,s_* spectrin bundles in our model generate a spring potential energy U*_s,S_* when they diverge from the resting length d*_o,S_* that is given by

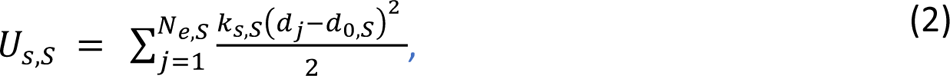

where k*_s,S_* is the spectrin constant and d*_j_* is the length of the edge *j* expanding between the *l* and *l*^’^nodes.

To simulate the evolution of the spectrin mesh to a relaxed configuration where the potential energy generated by its edges minimizes, we assumed, as in [27, 28], that the position ȓ_l_ ∈ ℝ^2^, *l* ∈ {1, …, N*_n,A_*} of the nodes representing the actin short filaments connecting the spectrin bundles freely moves at a velocity U¯_l_, and that adhesion complexes generate a viscous resistance to network deformations, given by ξU¯_l_, where ξ is a drag coefficient. This resistive force is balanced by the force generated by the spectrin potential energy at each node

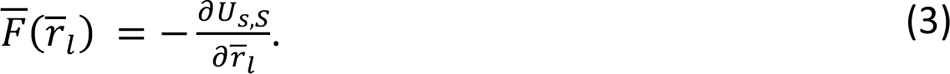

Thus, ȓ_l_ evolves according to

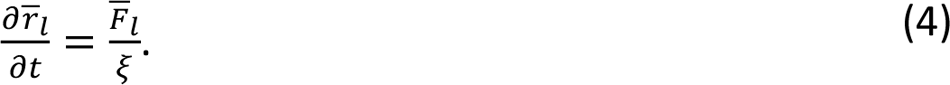

The simulations were run in MATLAB_R2021a on a desktop computer. We used the delaunayTriangulation.m function to initialize the spectrin triangular mesh. Then, we solved Eq. 4 with the Euler method using small time-steps Δ*t* to ensure numerical stability. We evolved the system until there was no significant change in its configuration, i.e., until the system was “equilibrated”. For Figure 6A the equilibration time is 60 seconds, for Figure 6D-E 120 seconds, and for Figure 6G-I 600 seconds. Since we were interested in the evolution between two different configurations of the spectrin mesh, instead of taking the mechanical parameters of the model from the literature related to one configuration or another, we fitted these parameters to qualitatively match the experimental observations. Other parameters, such as the length of the spectrin bundles, were taken from the literature. The units of the parameters were set to match the scales in the experiments. All the model parameters are in Table 3.

In Figure 6C, we introduced edges with a cable element in the model that produces a potential energy given by

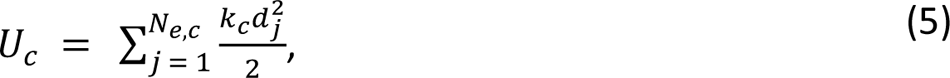

where k_c_ is the cable constant. Note that the cable elements generate a force that shrinks the corresponding edges. Since they connect to the spectrin network on one end and points representing focal adhesions with zero velocity to the other end, these cable elements stretch the spectrin network. Note that the evolution of the network is now given by the balance of all the forces acting in the cell cytoskeleton, i.e.,

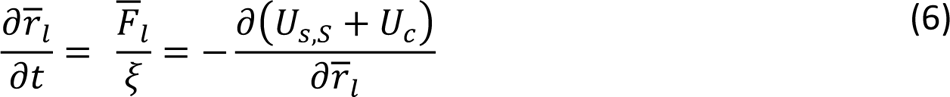

A recent study using molecular dynamics simulations [32] shows that an actin-spectrin model under a low strain rate is more prone to exhibit detachment of the actin-spectrin interphase rather than fragmentation of the spectrin bundle. We included this observation by detaching the edges corresponding to spectrin bundles when an expanding force generated by their spring potential U*_s,S_* is greater than a force threshold *F*_th_. We simulated the spectrin detachment from short actin filament by eliminating the corresponding edge from the network.

However, in the experimental data, the spectrin mesh was constrained by stress fibers. Therefore, we included stress fibers to the model by adding the corresponding edges to the top and the bottom of the spectrin mesh. As in [28], these edges have a spring and a cable element with constants k*_s,F_* and k_c,F_, respectively, creating a potential energy U_F_ = U_s,F_ + U_c,F_. The contraction generated by the cable element resembles the contraction generated by the myosin rods along the long actin filaments of the stress fibers. The stress fiber resting length d*_o,F_* corresponds to the calculated edge length in a relaxed state. Focal adhesions delimit stress fibers. The length of the edge connecting to focal adhesions was initially larger than the other edges in the stress fibers because we assumed that actin polymerization at the focal adhesions and actin exchange along the stress fiber also generate forces that affect the stress fibers resting state, as in [33]. Thus, this initial arrangement produces an instability that increases the length of all the stress fiber edges, thereby increasing their generated force and pulling the spectrin mesh to the focal adhesions. Moreover, during experiments, the stress fibers actively moved toward each other. To mimic this motion, the focal adhesion points moved vertically towards each other at a constant velocity U¯_A_ during the first half of the simulation (300 seconds) and at zero velocity afterward.

We introduced myosin linkers connecting the spectrin mesh to the stress fibers in the model. The myosin linkers initially span from an extreme node of the stress fibers to the center of the closest triangle formed by spectrin bundles in the spectrin mesh. These edges have a cable element with constant k*_c,L_*. Thus, they generate a shrinking force that affects the corresponding node in the stress fiber and the nodes of the spectrin triangle (the force is equally distributed among the three nodes of the spectrin triangle). We assumed that these myosin linkers do not shrink indefinitely, instead they detach from the network if their lengths are less than d*_min_*. When one of the sides of the spectrin triangle where a myosin linker connects detaches, then the myosin linker detaches from that spectrin triangle, and attaches to a randomly selected free spectrin triangle within a radius ȓ, d_min_ < ȓ < d*_max_*. If there are no free spectrin the myosin linker is removed from the network. We also added myosin rods inside the spectrin mesh. These rods attach to spectrin triangles in both extremes and have a cable element with constant k*_c,M_*. As the myosin linkers, if one of the sides of one spectrin triangle detaches, the corresponding extreme of the myosin rod randomly selects a free spectrin triangle within a radius ȓ, d*_min_* < ȓ < d*_max_*, to connect. If there are no free spectrin triangles within that radius, then the myosin rod is removed from the network. Furthermore, myosin rods are randomly created and removed from the spectrin mesh at a rate ф*_a_* and ф_r_, respectively. This was implemented in the simulation by adding (removing) a myosin rod at each time-step if a randomly drawn number from a uniform distribution *u*∼U(0,1) is less than the probability Δ*t*ф*_a(r)_*. These rates and the initial number of myosin rods were informed by experimental observations. The simulation of the full cytoskeleton network composed of edges representing spectrin bundles, stress fibers, myosin linkers or myosin rods, and nodes representing short actin filaments, focal adhesions, or stress fiber connectors, is described below:

At the start of the simulation:

- Initialize the network with the spectrin bundles, short actin filaments, stress fibers, focal adhesions, and myosin linkers.
- Add the myosin rods to the spectrin mesh to randomly selected locations, verifying that their length is within d*_min_* and d*_max_*.
- At each time-step:
  - For each *j* edge in the network spanning from ȓ*_l_* to ȓ*_l_*’:
  - Calculate the force generated by the potential energy of their spring and/or cable elements at 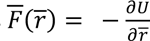 and ȓ*_l_*’.
  - Remove from the network an edge corresponding spectrin bundles if it generates a force larger than *F_th_*.

- Update the position of the nodes, by adding the force generated by their edges, 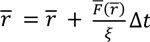
  - For the focal adhesions: If time < *t_s_*, move the nodes corresponding to the focal adhesions vertically at velocity U¯_A_, hence ȓ = ȓ + U¯_A_Δ*t*. Otherwise, set the velocity to zero.
  - For the edges corresponding to myosin (rods or linkers) attached to a spectrin triangle that lost one of its sides: find a new free spectrin triangle to attach within a radius d*_min_* < ȓ < d*_max_*. If there are no free spectrin triangles within that radius, remove the spectrin edge from the network.

- If *u* ∼ U(0,1) < Δ*t*ф_a_, add a myosin rod to a random location in the spectrin mesh.
- If *u* ∼ U(0,1) < Δ*t*ф*_r_*, randomly select a myosin rod and remove it from the network.

To link experimental observations of spectrin condensates in low tension state using FRET (Figure 5C-D) with our simulations, we plot the same network (Supplementary Figure 6D,E) but color-coded for tension instead of force. To calculate the tension, we divide the force generated by the spring element of each bundle over their length. Results in Supplementary Figure 6B-E show that locations with low force have low stress. Note that in the simulated network (Supplementary Figure S6B-E), most spectrin bundles have low force (tension) and those few bundles with high force (tension) are the result of the boundary conditions (stress fibers). Therefore, we conclude that the results of the simulated network are compatible with the low tension state observed in experiments.

## Figures and Figure Legends

**Supplementary Figure 1.**
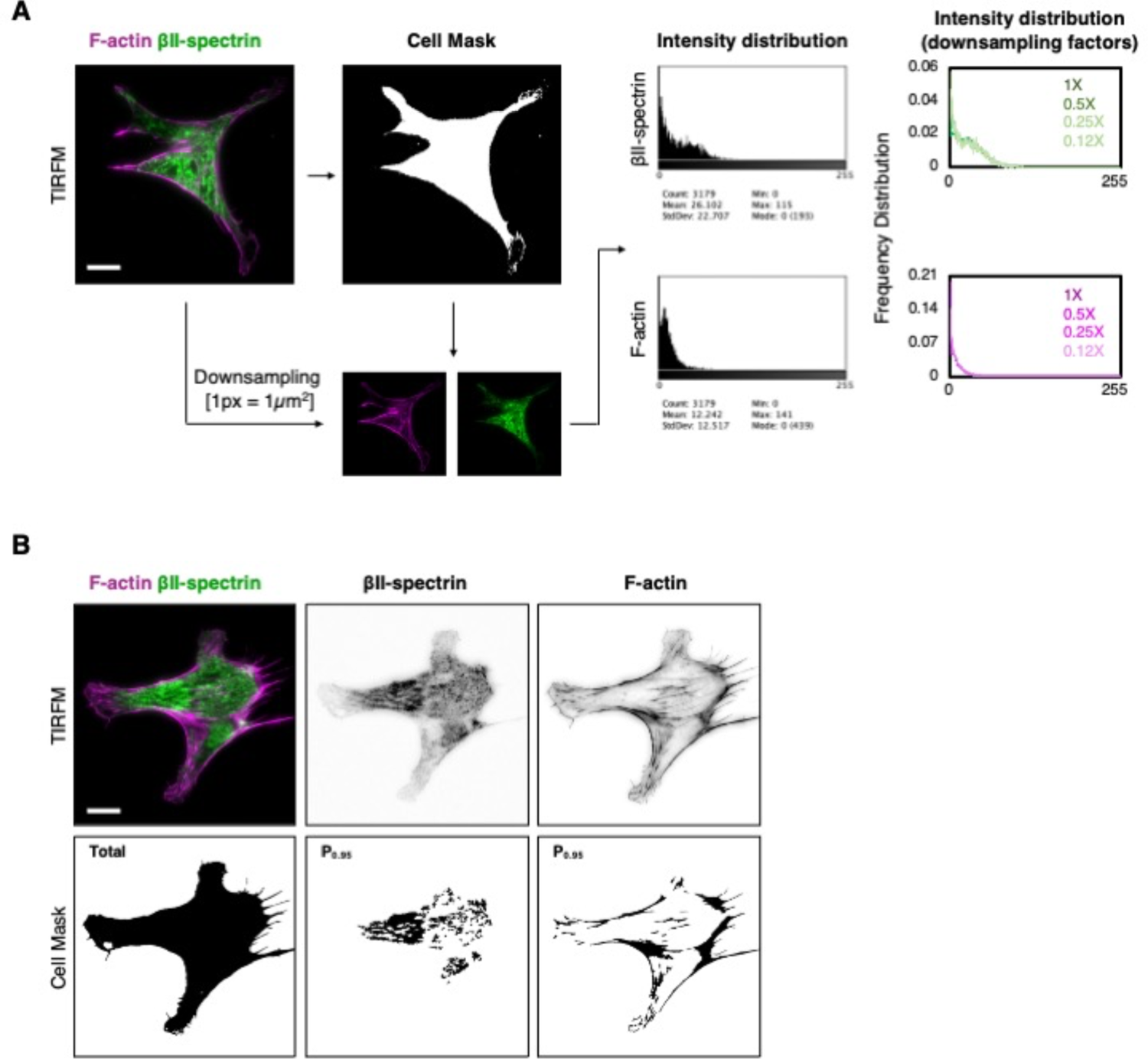
A) Experimental pipeline implemented to analyze the distribution of fluorescence intensities shown in Figure 1B (scale bar = 20 μm). Background-subtracted TIRFM images were used to obtain binary cell masks; single channel images were downscaled to 1 pixel = 1 μm^2^, with average interpolation. Intensity distributions for each single channel image was obtained within the cell mask, with the total count matching the cell area in µm^2^. Frequency distributions for n = 40 cells were merged and shown in Figure 1D. Fiji histogram output related to single-cell intensity distribution is shown. Different downscaling factors were applied on the cell shown; the pipeline here implemented did not introduce any bias. B) Representative analysis of P_0.95_ signal intensity. Left-tail threshold on background-subtracted and noise filtered images was consistently applied, only pixels with the 5% most intense signal were used to generate binary masks (P_0.95_ of total signal intensity distribution). Particle analysis was then performed to extract area and shape descriptor parameters. Representative images for βII-spectrin (green) and F-actin (phalloidin magenta) are shown, with corresponding P_0.95_ binary masks.

**Supplementary Figure 2.**
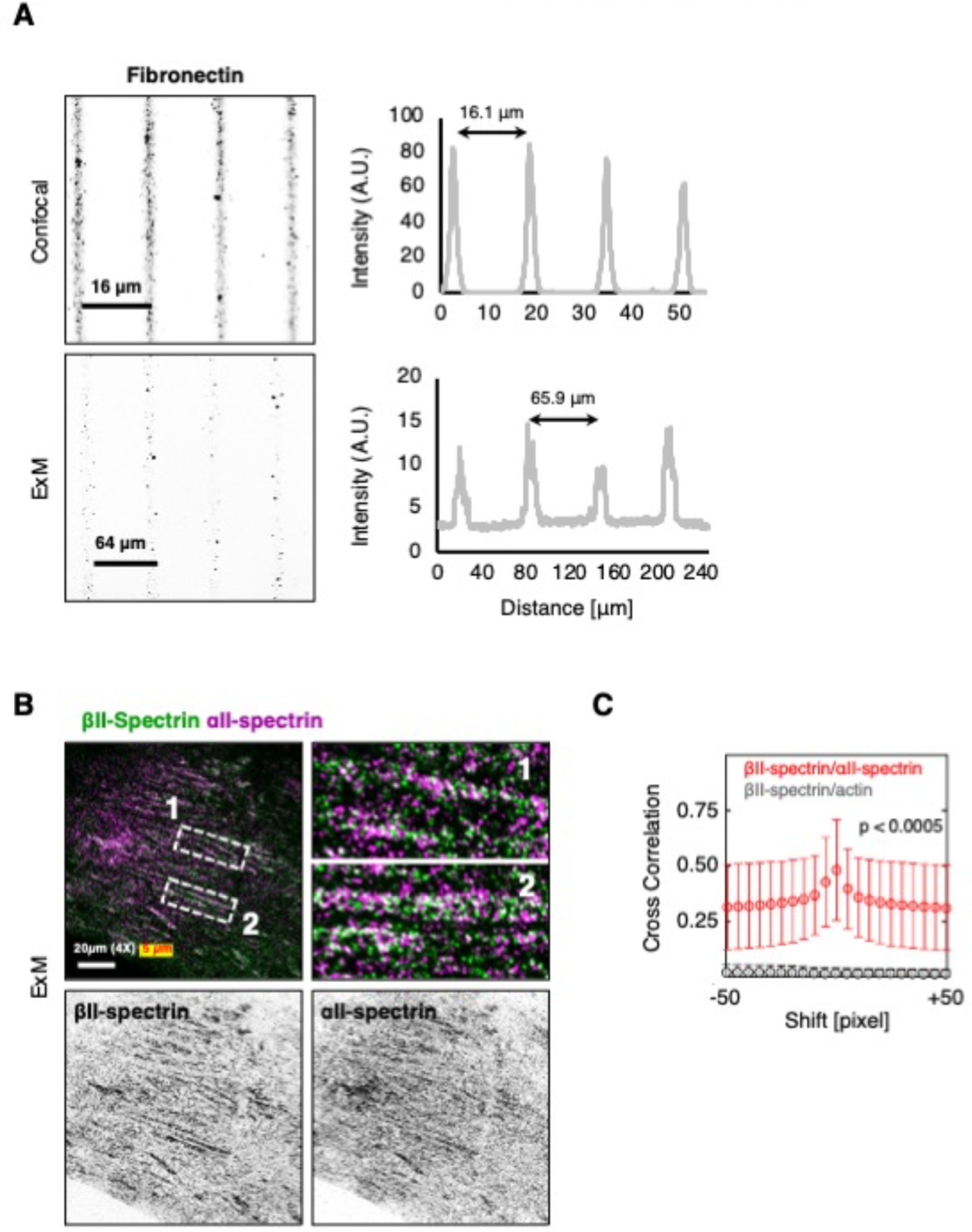
A) The estimated expansion factor for the ExM protocol was obtained by immunolabeling fibronectin-coated microprinted lines with an anti-fibronectin antibody, and imaged before and after expansion by confocal microscopy. The photolithography mask is designed with continuous lines of 4 μm thickness and 12 μm gaps. The profile plots are showing the expected 16.1 μm peak-to-peak distance (12 + 4 μm), and the resulting 65.9 μm after gelation and expansion: ∼4x expansion factor is therefore homogeneously achieved. B) Representative ExM image of MEF immunolabeled for βII-spectrin (green) and αII-spectrin (magenta) is shown (scale bar = 20 μm). Zooms (1-2, white dashed boxes) are shown to highlight the intermingled nature of the two epitopes. C) 2D cross correlation analysis of dual immunolabeled MEF imaged by ExM (window size 50 x 50 pixels): βII-spectrin/αII-spectrin (red) and βII-spectrin/β-Actin (gray, data are presented as mean ± SD, n = 7-9 cells, statistical analysis two-way ANOVA).

**Supplementary Figure 3.**
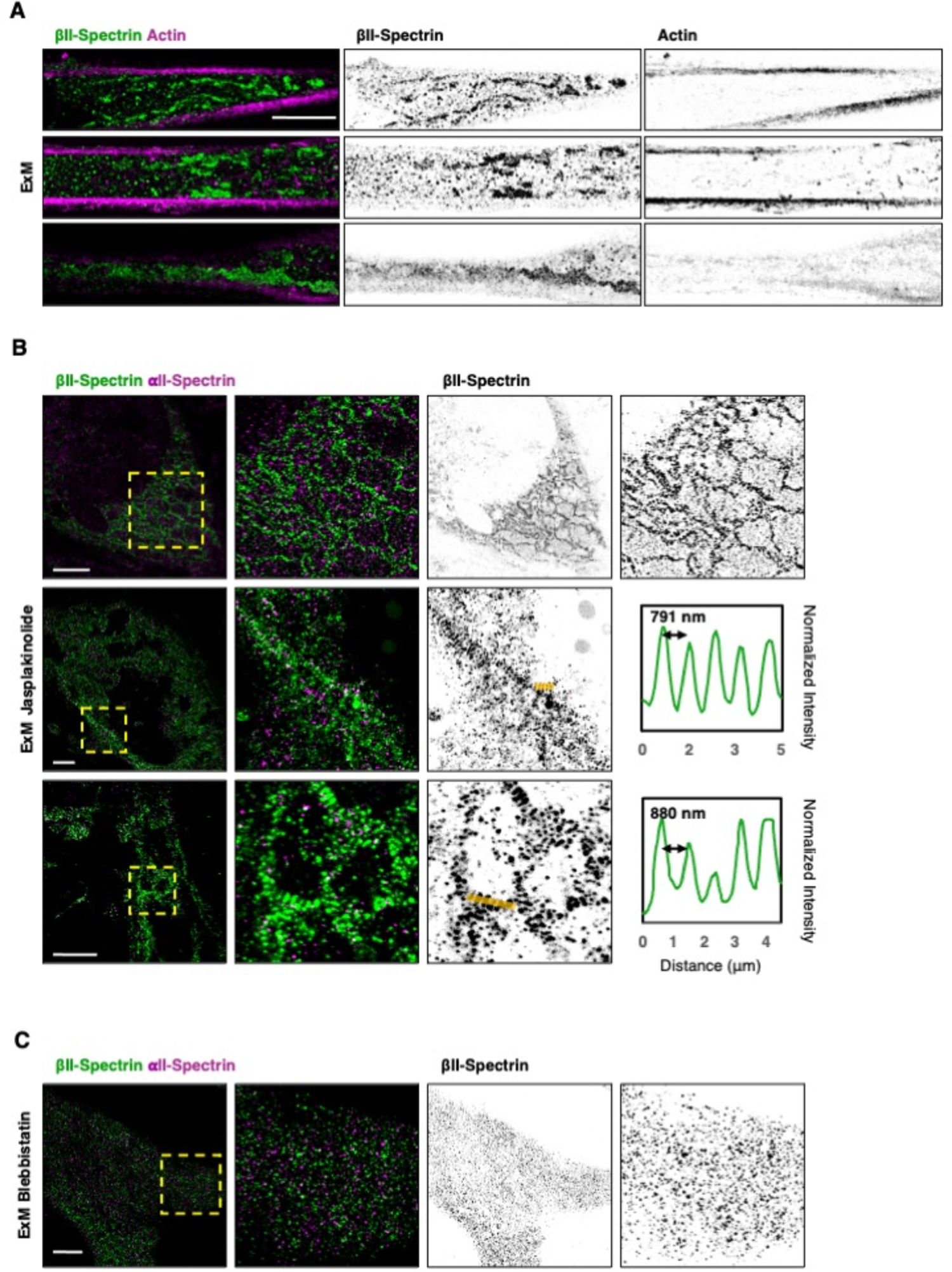
A) More representative ExM images of MEFs seeded on adherent microprinted pattern and immunolabeled for endogenous βII-spectrin (green) and actin (magenta). Similar representative images are shown also for Jasplakinolide (B) and Blebbistatin (C) treated MEFs (scale bars = 20 μm). Zooms are highlighted by the dashed yellow boxes. Intensity line-scans across the yellow rectangles are shown in the graphs and refer to the effect of Jasplakinolide treatment.

**Supplementary Figure 4.**
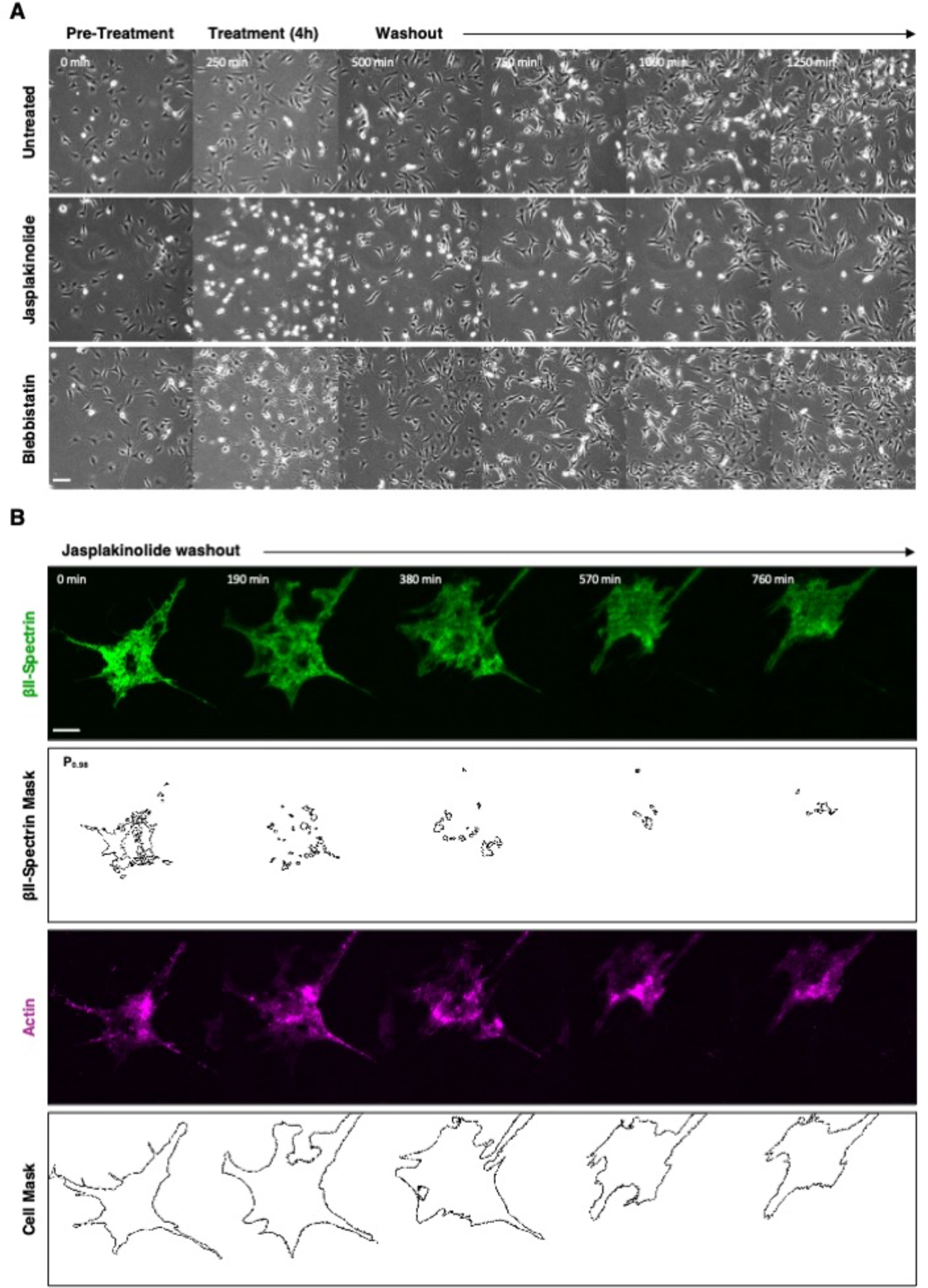
A) Phase contrast time lapse microscopy of MEF treated for 4 hours with Jasplakinolide 100 nM and Blebbistatin 10 μM. Cells were then followed for 16 hours after washout of the drugs to monitor recovery of cell shape and motility (scale bar = 100 μm). B) Live imaging by TIRF microscopy of MEF transiently transfected with GFP-βII-spectrin (green) and RFP-actin (magenta), treated with Jasplakinolide 100 nM. B) After 4 hours of treatment, cells were washed and imaged in fresh media for 14 hours by TIRFM, to monitor cell recovery from the treatment. Relevant frames are shown (scale bar = 20 μm). Outlines of the spectrin condensates (P_0.95_) over time, as well as cell mask based on the RFP-Actin signal, are shown to highlight recovery of cell motility and the evolution of spectrin condensates.

**Supplementary Figure 5.**
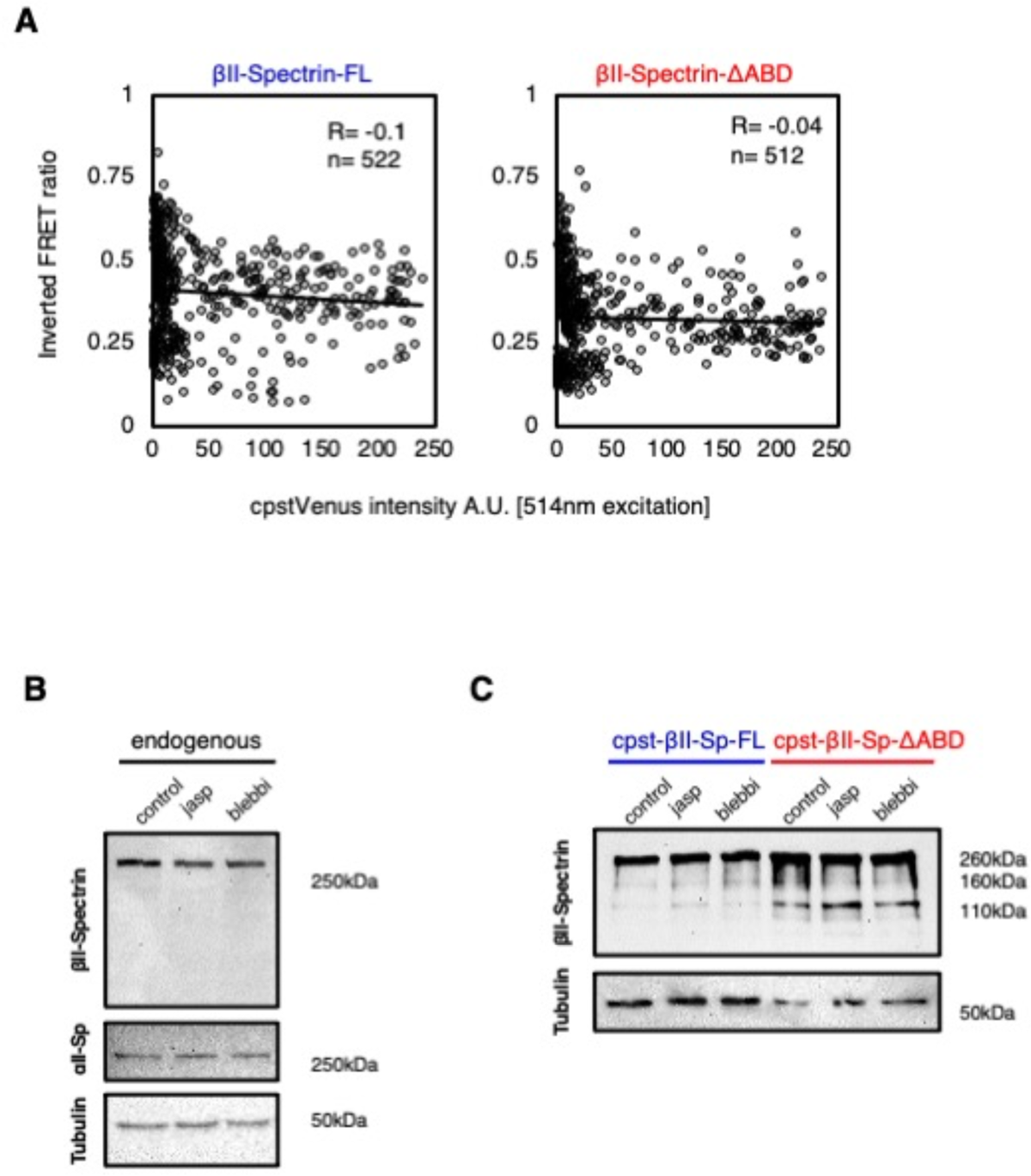
A) The graphs show the resulting whole-cell inverted FRET ratio as a function of the mean Venus signal intensity (excitation at 514nm) per each cell analyzed in this study (n = 522, 512). Pearson’s coefficients suggest no correlation exists between total fluorescent signal and the resulting FRET ratio. B-C) Western blot analysis of total cell lysates after 3-4 hours treatment with cytoskeletal impairing drugs implemented in this study. Immunoblots with anti βII-spectrin and anti αII-spectrin antibodies exclude protein degradation and fragmentation upon treatments. Anti-Tubulin immunoblot is used as loading control. The same analysis was performed on MEF transiently transfected with the two FRET-based constructs cpst-βII-spectrin-FL and ΔABD. Drug treatments did not produce differential degradation fragments compared to the control lane.

**Supplementary Figure 6.**
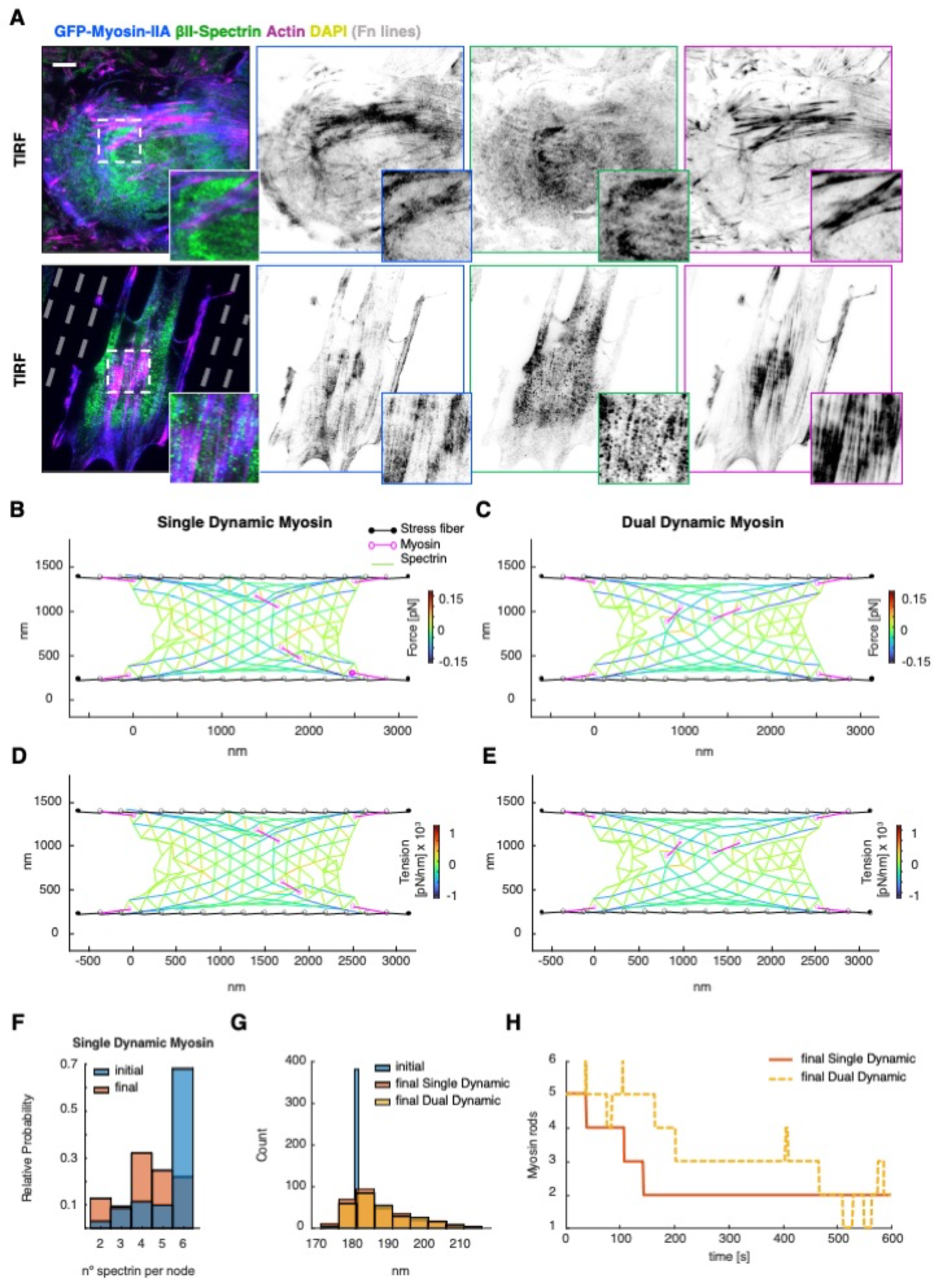
A) MEF transiently transfected with GFP-Myosin-IIA (blue), immunolabeled for βII-spectrin (green) and F-actin (phalloidin magenta), imaged by TIRFM (scale bar = 20 μm). Overlay and single channel images are shown, as well as zoom related to the white dashed box. The same microscopy analysis is performed on MEF seeded on microfabricated adhesive lines (4 μm adhesive cross-section, 12 μm non-adhesive surface) to force cell polarization and cytoskeletal orientation. Final configuration of the network in Figure 7F for single dynamic myosin (B) and double dynamic myosin (C), color-coded for the force generated by the spring element of the spectrin bundles (Eq. 3). D-E) Same as B-D but color-coded for tension. We calculated the tension dividing the force generated by the spring element of each bundle over their length. Length of the spectrin bundles in (F) and evolution of the myosin rods in the spectrin network over time (H), corresponding to the simulations in (B-C).

